# High-resolution mapping of the neutralizing and binding specificities of polyclonal rabbit serum elicited by HIV Env trimer immunization

**DOI:** 10.1101/2020.10.21.348623

**Authors:** Adam S. Dingens, Payal Pratap, Keara Malone, Sarah K. Hilton, Thomas Ketas, Christopher A. Cottrell, Julie Overbaugh, John P. Moore, P.J. Klasse, Andrew B. Ward, Jesse D. Bloom

## Abstract

Mapping the epitope specificities of polyclonal serum is critical to rational vaccine design. However, most high-resolution mapping approaches involve isolating and characterizing individual monoclonal antibodies, which incompletely defines the full polyclonal response. Here we use two complementary approaches to directly map the specificities of the neutralizing and binding antibodies of polyclonal anti-HIV-1 sera from rabbits immunized with BG505 Env SOSIP trimers. To map the neutralizing specificity, we used mutational antigenic profiling to determine how all amino-acid mutations in Env affected viral neutralization. To map the binding specificity, we used electron microscopy polyclonal epitope mapping (EMPEM) to directly visualize the Fabs in serum bound to Env trimers. Mutational antigenic profiling showed that the dominant neutralizing specificities were the C3/V5 and/or 241/289 glycan hole epitopes, which were generally only a subset of the more diverse binding specificities mapped with EMPEM. Additional differences between binding and neutralization reflected antigenicity differences between virus and soluble Env trimer. Further, mutational antigenic profiling was able to refine epitope specificity in residue-level detail directly from sera, revealing subtle differences across rabbits. Together, mutational antigenic profiling and EMPEM allow for a holistic view of the binding and neutralizing specificity of polyclonal sera and could be used to finely evaluate and guide vaccine design.

## Introduction

Mapping polyclonal antibody responses is central to understanding antigen-specific humoral immunity. However, it is difficult to disentangle the multiple epitope specificities within polyclonal responses. Often, serum neutralization assays or ELISAs with variant antigens are used to crudely map epitope specificities. But these and other traditional serological mapping approaches do not provide high-resolution, residue-level information for the multiple components of polyclonal serum responses. Cloning and characterizing many individual monoclonal antibodies (Scheid et al., 2009) has revolutionized our understanding of serum responses, but antibody cloning can be biased by the isolation strategy, is not proportional to antibody serum abundance or potency, and fails to characterize the entirety of the serum neutralization response. Further advances in understanding polyclonal sera have been made through techniques that rely on high-throughput B-cell receptor sequencing (Kreer et al., 2020), mass spectrometry-based approaches to directly sequence antibody proteins (Lavinder et al., 2014; Wine et al., 2013), or decomposing bulk serum-level measurements (Ackerman et al., 2017; Chung et al., 2015; Georgiev et al., 2013).

Only recently have techniques been developed that directly measure the antibody specificity in polyclonal sera. The first of these techniques, EMPEM, directly images serum Fabs bound to an antigen of interest (Barnes et al., 2020; Bianchi et al., 2018; Boyoglu-Barnum et al., 2020). However, this approach characterizes the binding response, whereas it is the neutralizing antibody response that is most directly correlated with vaccine protection. Here we combine EMPEM with a second complementary technique, mutational antigenic profiling (Dingens et al., 2017), that quantifies the effect of all single amino acid mutations to a viral entry protein on escape from serum neutralization.

For the purpose of this study, we mapped polyclonal anti-HIV antibody responses elicited with stabilized recombinant SOSIP Env trimers. These trimers have been used extensively as immunogens because they recapitulate the native or near-native structure of Env on the virus surface (Julien et al., 2013; Lyumkis et al., 2013; Pancera et al., 2014; Sanders et al., 2013; Sanders and Moore, 2017). In general, immunizing animals with prototypical SOSIP trimer variants based on the BG505 strain (Wu et al., 2006) induces autologous, tier-2 neutralizing antibody responses (De Taeye et al., 2015; Klasse et al., 2016; Sanders et al., 2015; Torrents de la Peña et al., 2018, 2017). While such immunizations can protect against infection of SHIV (Simian-Human Immunodeficiency Virus) bearing the matched BG505 Env in macaques (Pauthner et al., 2017, 2019), heterologous breadth has not been consistently achieved. This lack of breadth highlights the need to understand the targets of vaccine-elicited neutralizing antibodies and re-focus responses to more broadly conserved epitopes.

Prior mapping of SOSIP trimer-induced antibody responses in animal models has revealed viral strain- and species-specific hierarchical responses, with BG505 trimer immunogenicity in rabbits serving as a well-characterized model system. BG505-induced rabbit neutralizing antibodies predominantly target a BG505-specific glycan hole (GH) in Env’s glycan shield, centered on the broadly conserved glycosylation sites at residues 241 and 289 that are missing in BG505. Reintroducing glycans back in at these sites eliminates much of the neutralizing activity in many rabbit serum responses (Klasse et al., 2018, 2016) and mAbs isolated from immunized rabbits target this immunodominant glycan hole (McCoy et al., 2016). Serum neutralization assays with large panels of pseudovirus point mutants identified a second frequently immunogenic site in rabbits as the C3/V5 epitope (previously termed C3/465), as well as a less immunodominant and less commonly targeted epitope in V1 (Klasse et al., 2018). These rabbit immunogenicity data were largely corroborated using EMPEM to directly visualize serum Fabs bound to BG505 trimer bait (Bianchi et al., 2018). In SOSIP trimer-vaccinated guinea pigs and non-human primates, antibody cloning or EMPEM has identified additional strain-specific responses to the C3/V4, C3/V5, V1, and gp120/gp41 interface regions (Cottrell et al., 2020; Lei et al., 2019; Nogal et al., 2020, 2019). Additionally, trimer immunization often elicits non-neutralizing responses to the base of the trimer, a neo-epitope exposed on soluble trimers but inaccessible on viral membrane-bound Env (Bianchi et al., 2018; Cottrell et al., 2020; Hu et al., 2015; Kulp et al., 2017). Ongoing clinical trials will evaluate the immunogenicity of BG505 SOSIP trimer variants in humans (ClinicalTrials.gov Identifiers: NCT03699241, NCT04177355, and NCT03783130).

Below, we use mutational antigenic profiling to directly map the dominant neutralizing antibody specificities present within a panel of polyclonal sera from rabbits immunized the BG505 SOSIP trimer variants. In parallel, we map the binding specificities of these sera using EMPEM, giving us holistic view of both serum binding and neutralization.

## Results

### Rabbit sera panel

We chose a small panel of rabbit sera to optimize mutational antigenic profiling of polyclonal sera. We used sera from rabbits sequentially vaccinated with BG505 SOSIP trimer variants: either BG505 SOSIP.664, which contains the T332N mutation (Sanders et al., 2013) or the further stabilized BG505 SOSIP.V4.1 (De Taeye et al., 2015), each administered either 3 or 4 times. Details of the immunization schemes and characterization of some of these rabbits’ sera responses at earlier timepoints have been reported previously (Klasse et al., 2018, 2016; Ringe et al., 2019, 2017). We chose serum samples with various specificities, including sera that predominantly target the 241/289 GH or C3/V5 epitopes alone, both of these epitopes, or neither of these epitopes. To identify such sera, we performed preliminary TZM-bl neutralization assay mapping using pseudoviruses bearing mutations that affect each of these epitopes, as well as the V1 epitope rarely targeted in rabbits. The resulting sera panel and associated preliminary mapping data are shown in Figure 1A. These sera do not represent an unbiased collection of rabbit immune responses, but rather a curated selection of potent responses with different specificities.

**Figure 1.**
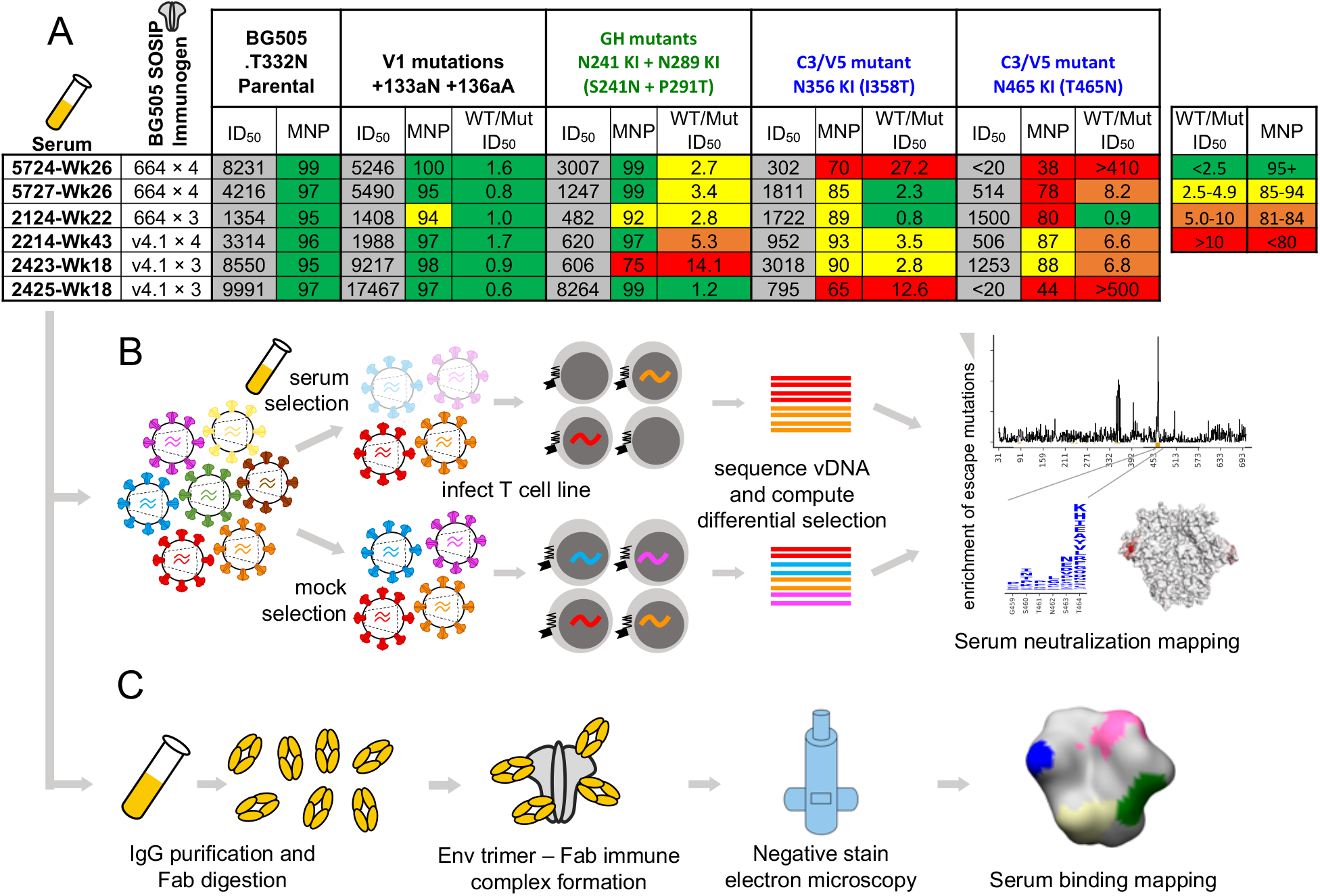
Overview of sera and epitope mapping approaches. **A.** Preliminary point-mutant mapping of the BG505-trimer vaccinated sera panel. The sera dilution that inhibits 50% of virus entry (ID_50_), maximum neutralization percentage plateau (MNP), and fold change in ID_50_ values of wildtype relative to mutant pseudoviruses (WT/Mut ID_50_) are shown for the parental BG505.T332N and pseudoviruses bearing insertion (V1 epitope) and glycan knock in mutation(s) (GH and C3/V5 epitopes). The weeks (Wk) post initial vaccination of the serum sample is specified in each sera’s name; sera are subsequently referred to by only their 4 digit ID number. **B.** Experimental schematic of mutational antigenic profiling. **C.** Experimental schematic of EMPEM.

### Mutational antigenic profiling of rabbit sera

We performed mutational antigenic profiling (Dingens et al., 2017) of each serum using libraries of replication-competent HIV virions expressing all mutants of the BG505.T332N Env (Haddox et al., 2018), allowing us to map autologous responses to the strain-matched BG505 trimer immunogen. Briefly, this approach (Figure 1B) first involves generating libraries of mutant viruses containing all single amino-acid mutations to Env compatible with viral replication. Each mutant virus library is then incubated with a highly selective concentration of sera before infecting a T-cell line such that only viruses that escape neutralization can enter cells. In our experiments, we chose serum concentrations to keep the average level of selection exerted by each serum relatively constant, averaging between 0.3% and 2.7% of the virions in the library escaping neutralization across replicates (Figure 1-figure supplement 1). The frequency of each mutation among viruses that are able to escape neutralization is quantified by Illumina sequencing of the viral cDNA produced in infected cells. Comparing the relative frequency of each mutation in the sera-selected condition to a non-selected control condition quantifies the effect of each mutation on resistance to sera neutralization (Figure 1B). As an additional control, we also incubated viral libraries with pre-vaccine sera for each rabbit. Figure 1-figure supplement 1 details the serum dilutions, the number of replicates (3 to 6 per post-immunization sera; median values are reported throughout), and the level of neutralization achieved by each serum in each experiment.

The Env mutations that affect neutralization by each serum are plotted in Figure 2. The results are largely concordant with prior knowledge on BG505 trimer immunogenicity in rabbits, with each serum targeting one or both of the C3/V5 or GH epitopes, which are indicated by blue and green, respectively, in Figure 2. Additionally, the antigenic profiling largely agrees with the crude epitope specificities mapped using a small panel of pseudovirus point mutants (Figure 1A, with mutations tested in preliminary TZM-bl assays plotted in black in Figure 2B). Notably, most sera select neutralization-escape mutations in the C3/V5 epitope to some extent, including sera from rabbit 2124, which appeared to predominantly target the GH based on the preliminary point mutant mapping (Figure 1A). Figure 2 is just one approach to visualizing these complex datasets; to facilitate more flexible data exploration, antigenic profiling data for each sera can be interactively explored in dms-view (Hilton et al., 2020) by visiting https://jbloomlab.github.io/Vacc_Rabbit_Sera_MAP/.This interactive visualization reduces potential interpretation and presentation biases by allowing users to examine the extent of neutralization escape by mutations at any site, and projecting the mutations onto interactive structures of the Env trimer or monomer.

**Figure 2.**
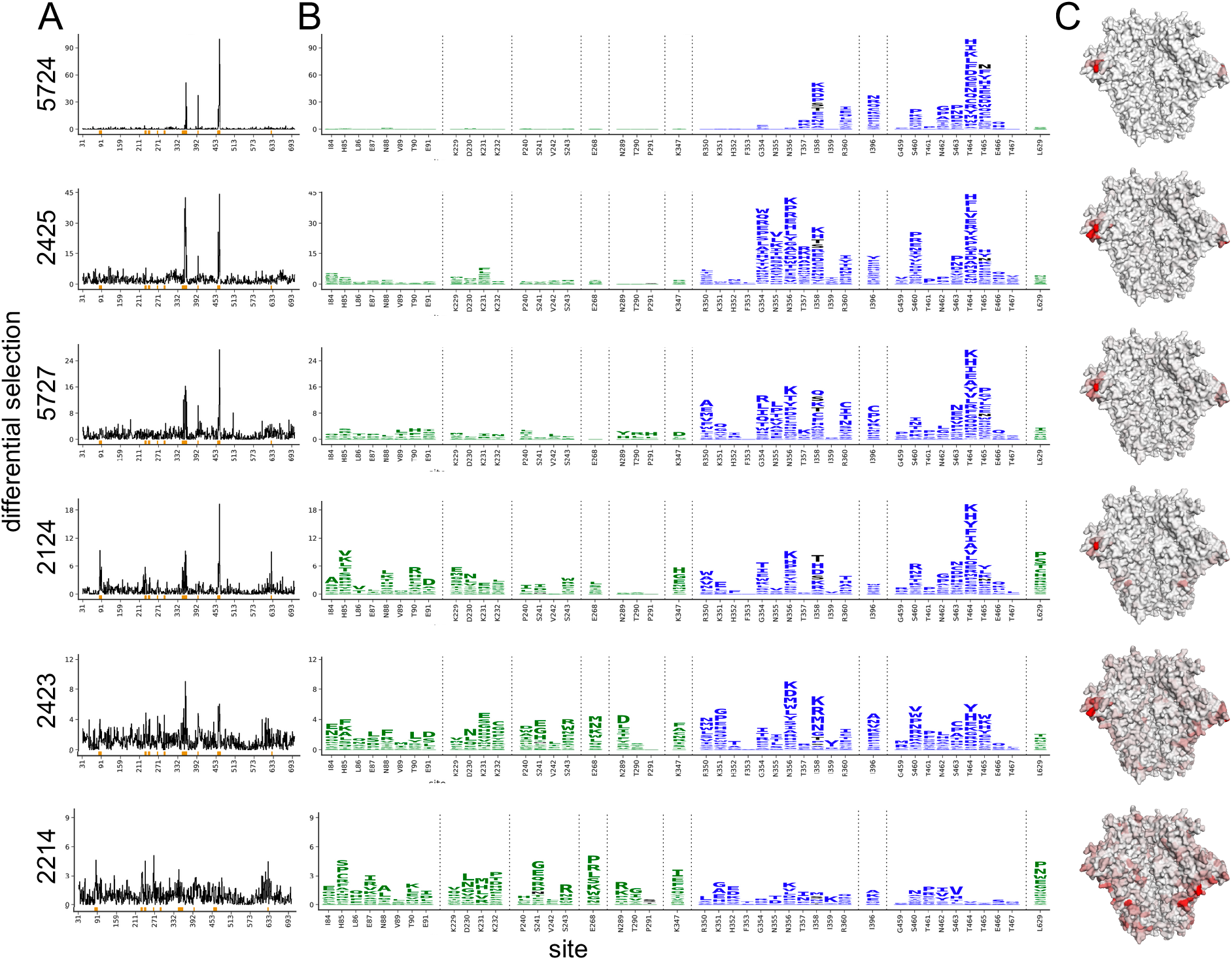
Serum neutralization-escape mutations mapped by mutational antigenic profiling. **A.** Line plots showing the positive site differential selection for each serum across the Env ectodomain. Differential selection is a measure of the enrichment of mutations in the sera-selected conditions relative to a no-sera control (see Methods for details). **B.** Differential selection for each mutation at key sites (indicated with orange underlines in A). The height of each letter is proportional to the differential selection for that amino-acid mutation. The GH epitope is colored green, and the C3/V5 epitope is colored blue. Mutations tested during preliminary point mutant mapping are colored black (Figure 1A; all mutations that add a tested glycan are indicated). For both A and B, The y-axis is scaled to the maximal effect size site for each sera; sera without a single dominant region of escape (2243 and 2214) are plotted such that 90% of the site-level signal is at <20% the y-axis maximum. **C.** The positive site differential selection is mapped onto the BG505 trimer (PDB: 5FYL). The color scheme is right censored at 15 to visualize subdominant responses; for samples with a maximum selection less than 15, the max value is mapped to most red color. An interactive version of these visualizations is at https://jbloomlab.github.io/Vacc_Rabbit_Sera_MAP/. Raw numerical values and logoplots for the entire Env ectodomain are in Figure 2–Source Data 1.

We hypothesized that sera with a few dominant sites of escape (e.g., serum 5724 in Figure 2) had neutralizing activity that was strongly focused on one epitope of Env, whereas sera with smaller-effect escape mutations to multiple regions (e.g., serum 2214 in Figure 2) had neutralizing responses that targeted multiple distinct epitopes. To explore this hypothesis, we tested if the small effect sizes observed in mutational antigenic profiling for some sera accurately reflect the effect of mutations in TZM-bl neutralization assays. We identified the most selected mutation at the most selected site for each serum, generated pseudoviruses bearing these mutations, and tested them in serum neutralization assays. The fold enrichment in mutational antigenic profiling was well correlated with the fold change in ID_50_ in the neutralization assays (Figure 2-figure supplement 7, Pearson’s r = 0.97). For example, the 5724 escape profile is focused entirely on the C3/V5 epitope: T464H is enriched ~190-fold in this serum’s mutational antigenic profiling and shifts the ID_50_ 64-fold in a TZM-bl neutralization assay (Figure 2-figure supplement 7). In contrast, 2423 targets both the C3/V5 and GH epitopes (Figure 2), and the maximal effect mutant N356K has just a ≈2-fold effect in both mutational antigenic profiling and neutralization assays (Figure 2-figure supplement 7). A caveat is that the extent of mutant enrichment upon serum selection is also influenced by the serum dilution used in experiments, as shown in Figure 2-figure supplement 1-6. However, the good correlation of the extent of focusing in the escape-mutation mapping and the TZM-bl neutralization assays suggests the mutational antigenic profiling data captures the amount of focusing in the neutralization response reasonably well.

### Contrasting the neutralizing and binding specificities of the polyclonal sera

To contrast the serum neutralization specificities described above with the serum binding specificities, we performed EMPEM on the same set of sera to directly visualize antibody binding to Env (Figure 1C). We reasoned that collecting both types of data would enable us to compare and contrast the sites where antibodies bind to the sites where mutations mediate escape from neutralization. Figure 3 presents the refined 3D reconstructions from negative stain electron microscopy of serum Fabs bound to immunogen-matched BG505 SOSIP trimer alongside mutational antigenic profiling data. Across all sera, it is immediately clear that the binding responses identified by EMPEM include many epitopes where mutations do not affect viral neutralization. For 5 of 6 sera, binding responses to both the GH and C3/V5 epitopes are observed. All sera contain additional binding responses to other epitopes. For example, even 5724, where we observe narrow, strongly focused viral escape in just the C3/V5 epitope, contains numerous additional binding responses, including the GH, N611 glycan, base-of-trimer, and V1/V3 epitopes.

**Figure 3.**
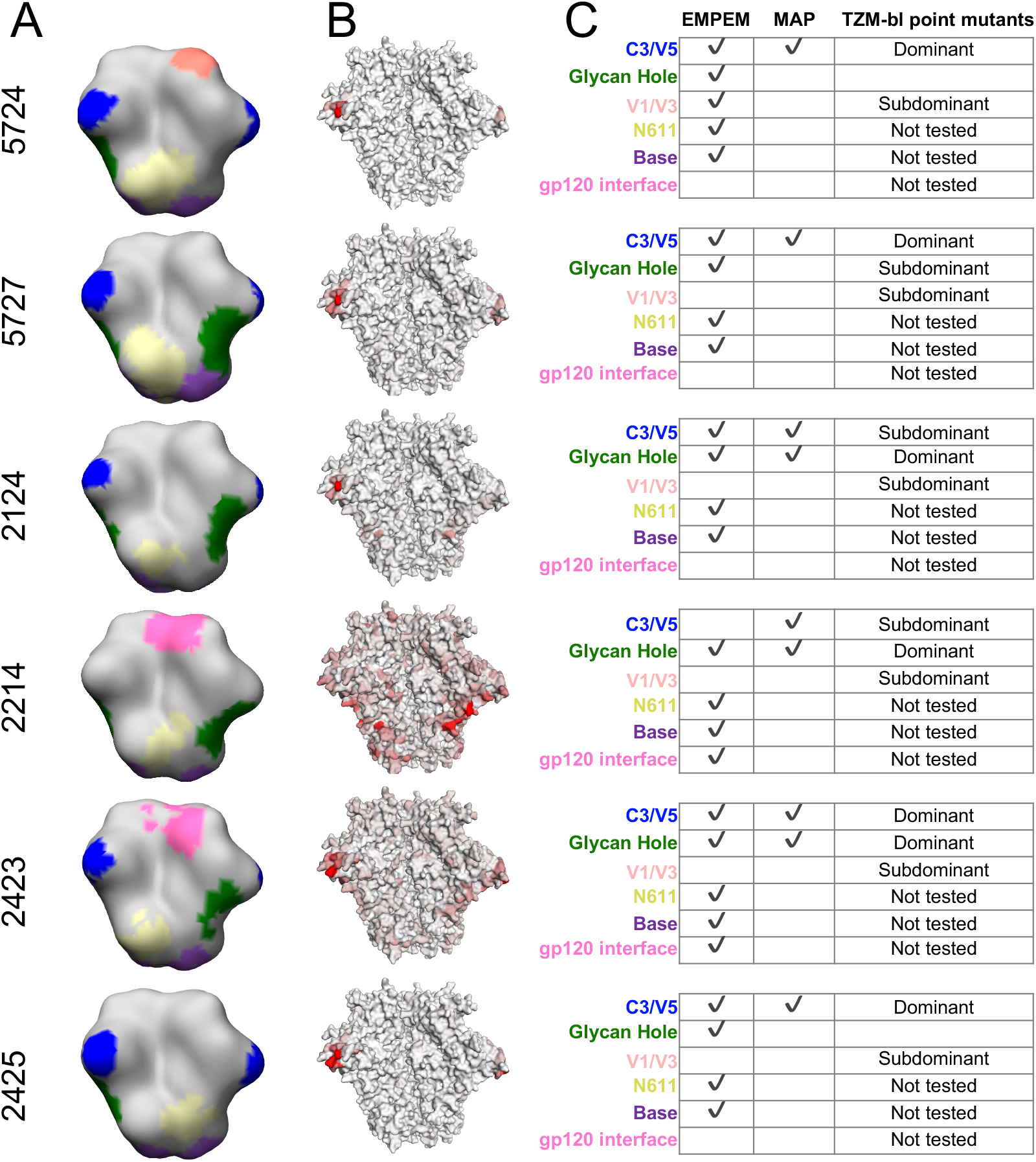
Comparing the binding and neutralization specificities of the sera. **A.** Refined 3D reconstructions from negative stain EMPEM. Specificities are mapped onto the Env trimer structure, with epitopes colored as in 3C. **B.** Mutational antigenic profiling data mapped onto the BG505 SOSIP Env structure, as in Figure 2C, represented here to contrast with EMPEM. **C.** Summary data from A and B, as well as validation TZM-bl neutralization assay point mutant mapping of single and double epitope knock out mutations (see Figure 3-Figure supplement 1 for details). We did not perform neutralization assay validation at epitopes where the discrepancies between the EMPEM and mutational antigenic profiling are easily explained by antigenicity differences between soluble trimer and virus (N611, base, and gp120 interface; labeled as “Not tested”).

Some of the differences in binding vs neutralization-escape are easily explained by antigenicity differences between stabilized Env trimers and replication competent virus. First, while the base-of-trimer epitope is presented on recombinant trimers and commonly elicited by trimer immunization (Bianchi et al., 2018; Cottrell et al., 2020; Hu et al., 2015), it is inaccessible in the replication-competent virus and hence does not elicit neutralizing antibodies. Therefore, base-binding responses are observed in all sera, but of course are not mapped as neutralizing epitopes. Second, binding responses to the N611 glycan region are also observed in all sera (Figure 3), but we do not observe viral escape by disrupting this glycosylation motif (Figure 2). It has been previously shown that site N611 is less glycosylated in BG505 SOSIP trimers than in virus, which enhances the immunogenicity of the exposed region when this glycan is missing (Derking et al., 2020). Accordingly, EMPEM identified binding responses to this region using the immunogen-matched trimer bait, while neutralizing responses are not apparent in mutational antigenic profiling because the virus libraries are likely to have higher glycan occupancy at N611. Third, gp120 interface responses are observed in two sera; both of these rabbits were immunized with BG505 SOSIP.v4 trimers. This gp120 interface epitope includes the A316W stabilizing mutation added to the SOSIP.v4 to reduce the exposure of the V3 loop (De Taeye et al., 2015). We have recently found that the A316W mutation alters immunogenicity to this region, eliciting mutation-specific responses to this region that would not cross-react with the A316-bearing viral libraries (manuscript in prep). Together, these trimer-binding but non-neutralizing responses to the base, N611 glycan, and gp120 interface reflect and expand on previously characterized antigenic differences between Env trimers to Env on the surface of virus (Bianchi et al., 2018; Cottrell et al., 2020; Derking et al., 2020; Hu et al., 2015).

The few remaining differences in binding and neutralization could reflect a number of factors. Some binding responses may be non-neutralizing, although it is often assumed that antibodies that bind native Env present on the virus – mimicked by stabilized Env trimers – are neutralizing (Burton et al., 2000; Yang et al., 2006). Alternatively, at the serum concentrations we tested, the antibody occupancy on the virions or the binding kinetics may disfavor neutralization. Further, some observed binding responses may be neutralizing but at such subdominant levels that they do not exert selective pressure in the mutational antigenic profiling at the serum concentrations tested. Alternatively, some epitopes may require saturation in order to be neutralizing and did not reach these saturating levels at the serum concentrations tested in mutational antigenic profiling.

To investigate these hypotheses, we tested each sera’s ability to neutralize pseudovirus point mutants bearing mutations to one or both of the C3/V5 and GH epitopes (Figure 3 – figure supplement 1 and summarized in Figure 3C). Comparing the effect of single epitope mutations to multiple epitope mutations can help define the relative dominance of different neutralizing specificities (Klasse et al., 2018). A caveat is that neutralization assays for different sets of mutants were performed in two different laboratories; we therefore only present general interpretations in Figure 3C. For both 5724 and 2425, single or multiple glycan knock in mutations to the GH epitope had negligible effects, while single or multiple glycan knock in mutations to the C3/V5 epitope had large effects (Figure 3 – figure supplement 1). For these two sera, knocking in a glycan to each epitope (S241N + I358T) did not have much larger effects than the single C3 glycan knock in (I358T), suggesting these sera have limited GH neutralizing responses even after eliminating some of the C3/V5 directed neutralizing response. While this matches the mutational antigenic profiling, binding GH responses are observed in these sera (Figure 3A). This suggests that these GH responses are either non-neutralizing or *much* less dominant than the C3/V5 neutralizing antibody responses. Of note, while the resolution of the negative stain EMPEM in this present study limits fine epitope interpretations, GH binding responses (specifically “GH2”-like responses) have previously been identified in non-neutralizing sera using cryo-EMPEM (Bianchi et al., 2018).

When the effect of a mutation to one (e.g. epitope A) is apparent when another epitope (e.g., epitope B) is knocked out but not when testing the epitope A mutation alone, we can interpret the neutralizing antibody response to epitope A as being “subdominant” relative to the epitope B response. Here, 5727, 2124, 2214, and 2423 all displayed a greater effect for the double epitope glycan knock in mutations (S241N + I358T) than either of the single epitope knock in mutations. Comparing the effects of the single and double epitope mutations suggests sera 2423 had relatively equivalent neutralizing responses to both the GH and C3/V5 epitopes, sera 2124 and 2214 had a dominant response to GH and a subdominant response to C3/V5, and sera 5727 had a dominant response to C3/V5 and a subdominant response to GH (Figure 3, Figure 3 – figure supplement 1). EMPEM identified binding responses to both epitopes in 3 of 4 of these sera, while mutational antigenic profiling identified neutralizing responses to both epitopes in only 2 of 4 (Figure 3).

Since there were a number of instances in which subdominant neutralizing responses to the GH or C3/V5 epitope identified with TZM-bl point mutant mapping did not appear in either EMPEM or mutational antigenic profiling (e.g. GH response for 5727 is absent in mutational antigenic profiling, and a C3/V5 response for 2214 is absent in EMPEM), we examined if there were additional unobserved subdominant responses. We focused on the effect of insertion mutations to V1, an epitope occasionally targeted in rabbits (Klasse et al., 2018). We tested sera for neutralization of V1 and V1 + GH epitope mutants and compared effects to GH mutations alone. While V1 insertions alone did not affect neutralization of any sera, they had an additional effect when tested with GH mutations relative to GH mutations alone (Figure 3 – Figure supplement 1). This suggests that there are subdominant neutralizing antibody responses to V1 in all sera – though we cannot rule out that the V1 + GH double mutants broadly affect antigenicity. Mutational antigenic profiling did not clearly identify any V1 responses, while EMPEM identified a V1/V3 binding response – that overlaps the V1 insertion mutations – in one serum (5724, Figure 3).

### Residue-level refinement of sera epitopes

The mutational antigenic profiling data also allows for residue-level refinement of the dominant targets of the neutralizing antibody response. For example, it is immediately apparent that the clustered, surface-exposed sites 464 and 356 and/or 358 “anchor” the C3/V5 epitope, with many mutations at these sites having large effects for nearly all sera that strongly target this epitope (Figure 4). However, detailed epitope specificity at other C3/V5 sites varies across rabbits: some sera are more focused on various regions of C3, including sites 350 and 351 (e.g., sera 5727, 2423, 2124), or 354-357 (e.g., sera 5727, 2423, 2124, 2425), whereas serum 5724 is more narrowly focused on the V5 region previously identified by knocking in a glycan at site 465. Notably, site 396 bridges these two epitope regions in both linear sequence and structural space and is a site of escape for most sera.

**Figure 4.**
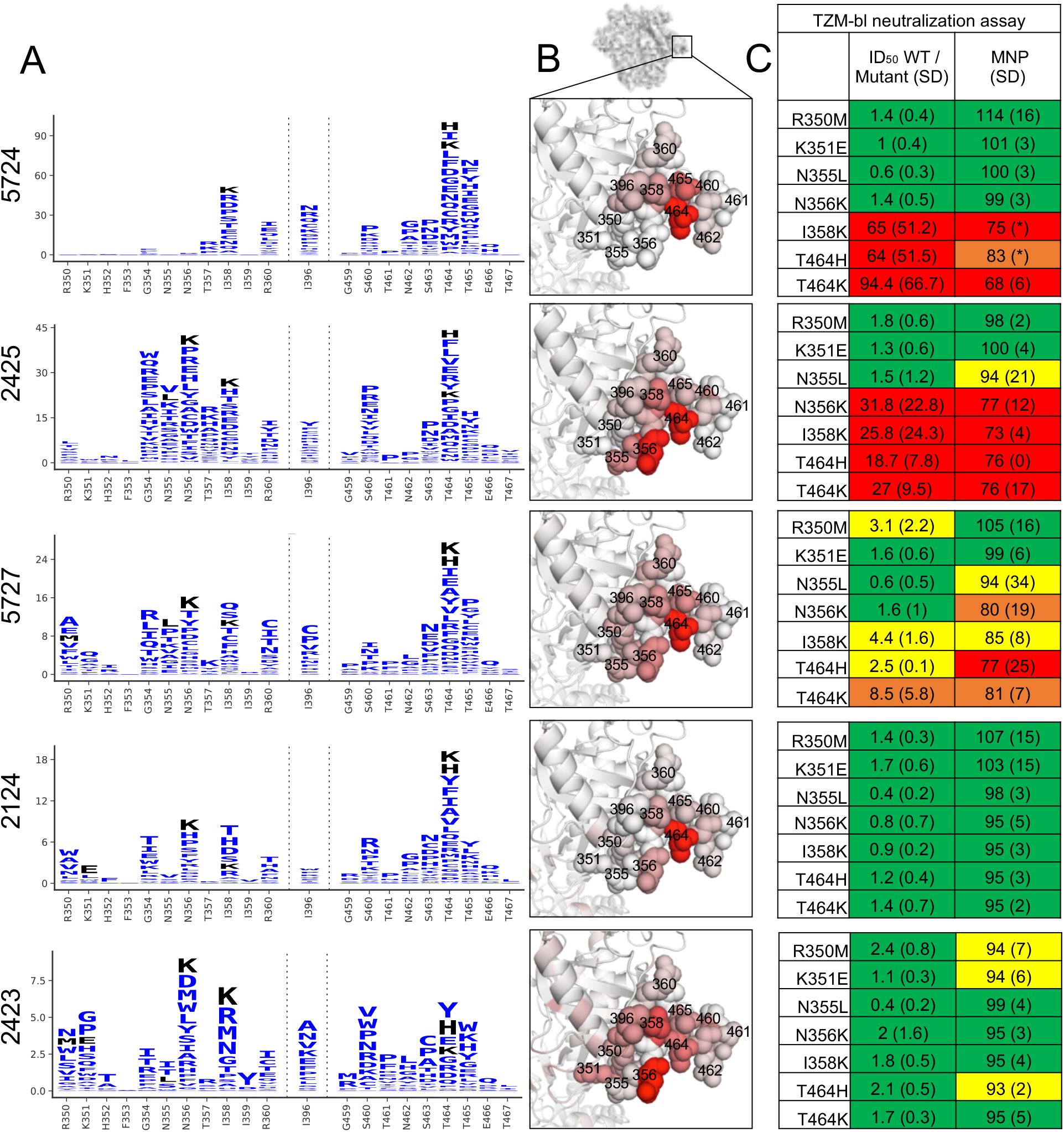
C3/V5 residue-level epitope specificity. **A.** Differential selection is plotted in logo plots for the C3/V5 epitope, as in Figure 1b. Mutations validated in TZM-bl neutralization assays (C) are colored black. The Y axis is scaled to the largest effect size site. **B**. The positive site differential selection is mapped onto the BG505 trimer (PDB: 5FYL), colored as in Figure 1C. Sites shown in A are shown with spheres in B. These data can be explored interactively using dms-view at https://jbloomlab.github.io/Vacc_Rabbit_Sera_MAP/. **C**. The fold change in ID_50_ relative to wildtype and the maximum neutralization plateau (MNP) for each mutant validated in a pseudovirus TZM-bl neutralization assay. The table color scheme is as in Figure 1A, and the standard deviation of replicate measures are shown in parentheses. See Figure 4-Figure supplement 1 for the single sera with relatively less targeting of this epitope.

We validated this residue-level specificity using TZM-bl neutralization assays (Figure 4). For example, mutations to sites 350, 351, 355, and 356 have very little effect on 5724, while mutations to 358 and 464 have large effects. Similarly, 2425 is most focused on residues 354-358 in the C3 region, mutations to these sites gave larger effects while those to 350 and 251 did not. While the magnitude of effect size varied across sera (note the differing y axis in Figure 4), the TZM-bl mapping generally reflected these effect sizes.

It is also clear that the mutational antigenic profiling better explores the effects of different possible escape mutations than testing smaller panels of psuedovirus mutants. For example, while knocking in a glycosylation site at site 465 (T465N) escapes many of the sera as previously shown (Klasse et al., 2018), other mutants at this site have similar or even larger effects (Figure 2, 3). This suggests that the immune response is directed at site 465 and neighboring V5 residues, as opposed to targeting this general epitope region that is obstructed by adding a bulky glycan to site 465.

There is greater variation in both the residue-level specificity and magnitude of neutralizing antibody responses to the glycan hole epitope. However, overlaying data onto the trimer structures make it clear that a subset of the sera target the 241/289 glycan hole. Examining data for three sera with clear enrichment in this region reveal selective pressure in a large epitope region, spanning from site 85 near the fusion peptide, through the 241/289 GH region, and extending to site 347 in/near the C3/V5 epitope. While we arbitrarily classify site 347 as part of the glycan hole epitope because enrichment at this site more closely tracks with sera that target the glycan hole epitope (Figure 2), the blending of these epitopes supports the notion that the C3/V5 and GH epitopes together constitute a BG505-specific immunodominant “super-epitope” in rabbits (Nogal et al., 2020). Mutations at site 629, particularly L629P, are also enriched strongly in two sera that target the GH epitope (Figure 5). Site 629 is near the N-terminus of the HR-2, located below and buried beneath the GH epitope in the pre-fusion structure; the proline mutation may disrupt the HR-2 alpha helix, altering its antigenicity or accessibility of the GH epitope.

**Figure 5.**
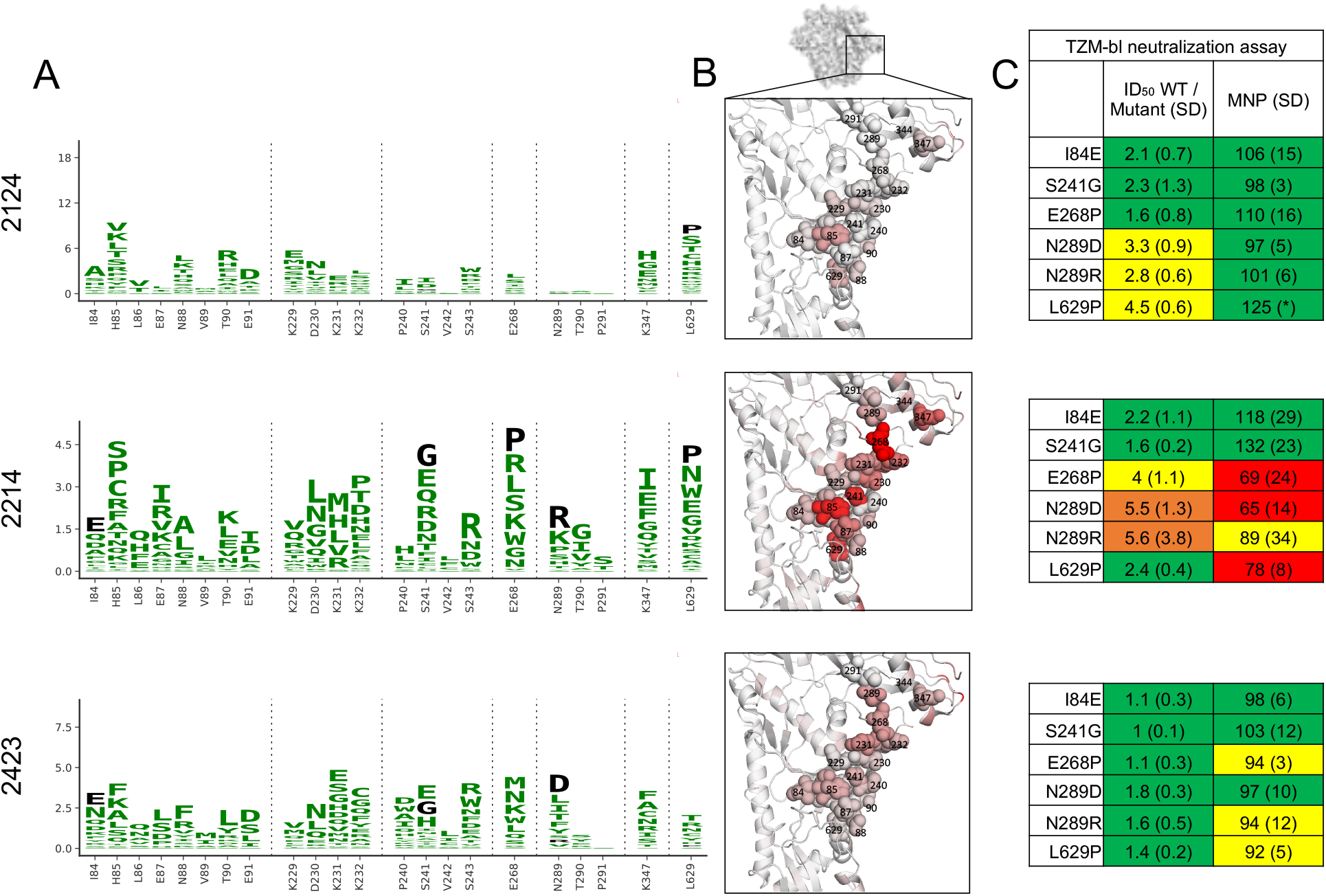
Glycan hole epitope specificity. **A.** Differential selection is plotted in logo plots for the glycan hole epitope, as in Figure 1b. Mutations validated in TZM-bl neutralization assays (C) are colored black. The Y axis is scaled to the largest effect size site, which is oftentimes not in this GH view. **B**. The positive site differential selection is mapped onto the BG505 trimer (PDB: 5FYL), with a single monomer shown and colored as in Figure 1C. Sites shown in A are shown with spheres in B. These data can be explored interactively using dms-view at https://jbloomlab.github.io/Vacc_Rabbit_Sera_MAP/. **C**. The fold change in ID_50_ relative to wildtype and the maximum neutralization plateau (MNP) for each mutant validated in a pseudovirus TZM-bl neutralization assay. The table color scheme is as in Figure 1A, and the standard deviation of replicate measures are shown in parentheses. The table color scheme is as in Figure 1A. See Figure 5-Figure supplement 1 for the three sera with limited targeting of this epitope.

Mutations in the GH epitope were oftentimes only moderately enriched compared to the C3/V5 epitope, and TZM-bl assays also showed only small shifts in the neutralization curve for these mutants. Sera without clear enrichment of escape mutations in this epitope also were not affected by epitope mutations in TZM-bl neutralization assays (Figure 5-Figure supplement 1).

## Discussion

We have used mutational antigenic profiling to map the dominant neutralizing antibody specificities in polyclonal rabbit sera elicited with Env trimer immunization. In parallel, we used EMPEM to map the total serum binding specificities. Contrasting the serum binding and neutralizing specificities suggests that the dominant neutralizing antibody responses are only a subset of binding responses – even when just examining bona fide neutralizing antibody epitopes. Additional differences in binding and neutralization highlight antigenicity differences between soluble SOSIP Env trimers and Env on the surface of viruses, which is relevant to the use of trimers as immunogens.

This work also shows the utility of mutational antigenic profiling in mapping polyclonal serum responses to HIV Env. We recapitulate and extend prior knowledge on rabbit antibody responses to BG505 trimer immunization (Bianchi et al., 2018; Klasse et al., 2018; McCoy et al., 2016), with the C3/V5 and glycan hole regions being the dominant immunogenic epitopes. We map responses to multiple epitopes of polyclonal serum responses at once, a significant advance over more traditional mapping approaches. Further, we refine the residue-level specificity of these epitopes directly from sera, revealing fine-grain epitope differences. For example, some C3/V5 responses were more focused on the C3 region than others. But there were sites that appeared to “anchor” this epitope – all sera targeting this epitope were strongly affected by mutations to sites 358 and 454. It remains to be determined if these different epitope specificities are the result of monoclonal responses with varying binding footprints, or if they vary due to differing sets of multiple overlapping antibodies targeting the same epitope region.

The glycan hole epitope it is not a desirable broadly neutralizing epitope that can be exploited for vaccine design (Yang et al., 2020), and the C3/V5 epitope is similarly problematic due to its relatively low sequence conservation. While the autologous neutralizing responses we mapped here are encouraging, we hypothesize it will be important to silence these immunodominant and autologous neutralizing responses to aid in redirecting response to more broadly conserved epitopes. Our mutational antigenic profiling provides a rich map of potential alterations that disrupt these epitopes, which could be of use in resurfacing trimer immunogens.

While this work has allowed us to compare total polyclonal serum neutralization and binding, there are important limitations to keep in mind. One key point is that these methods have different sensitivities for dominant versus subdominant responses, making it impossible to directly quantify serum neutralization and binding on the same scale. Indeed, TZM-bl neutralization assays using single and double epitope mutant pseudoviruses suggests that both approaches may have “missed” subdominant neutralizing antibody responses to the V1 epitopes (Figure 3 and Figure 3 – figure supplement 1). Further rigorous quantification of responses – and comparisons of responses across sera – are not yet possible with existing technology and analytical approaches. Further, while mutational antigenic profiling examines the entire serum neutralizing antibody response, EMPEM maps the binding of IgG Fabs after purification. Lastly, easily visualizing and interpreting these complex datasets remains difficult, a challenge we have attempted to overcome here with the interactive visualizations available at https://jbloomlab.github.io/Vacc_Rabbit_Sera_MAP/. Nonetheless, this work describes the most detailed mapping yet of the specificities of polyclonal sera at both the binding and neutralizing level.

While both rational, structure-based vaccine design (Alam et al., 2017; Correia et al., 2014; Dubrovskaya et al., 2019; Haynes et al., 2012; Kong et al., 2019; Kwong and Mascola, 2018; Saunders et al., 2017; Xu et al., 2018) and broadly neutralizing antibody germline-targeting (Briney et al., 2016; Dosenovic et al., 2015; Escolano et al., 2019, 2016; Jardine et al., 2013, 2016; Medina-Ramírez et al., 2017; Steichen et al., 2019, 2016; Tian et al., 2016) approaches have begun to show exciting promise for HIV, continued progress will require iterative rounds of evaluating vaccine responses and redesigning immunogens and vaccine regimens (Ward and Wilson, 2020). Determining the extent of immunofocusing to targeted epitopes, while also tracking – and subsequently eliminating – off-target responses to less conserved regions will be critical to these efforts. Together, EMPEM and mutational antigenic profiling can aid in rational vaccine design by directly mapping polyclonal serum specificities for both binding and neutralization.

## Methods

### Mutational antigenic profiling

Mutational antigenic profiling was performed as previously described (Dingens et al., 2017). Briefly, 5×10^5^ infectious units of BG505.T332N mutant virus libraries (Haddox et al., 2018) were neutralized with serum dilutions for 1 hour at 37C as specified in Figure 1-Figure supplement 1. We performed additional experimental replicates for sera with smaller effect sizes in order to increase our signal relative to noise (Figure 1-figure supplement 1). Different library numbers (e.g. 1, 2, 3) are viral libraries generated from independently generated DNA libraries (Haddox et al., 2018), and letter labels (e.g. a, b, c) correspond to experiments done on different days with matched non-selected control library selections. After neutralization, viral libraries were then infected into 1×10^6^ SupT1.CCR5 cells in R10 containing 100μg/mL DEAE-dextran. Three hours post infection, the cells were resuspended in one mL R10 (RPMI [GE Healthcare Life Sciences; SH30255.01], supplemented with 10% FBS, 1% 200 mM L-glutamine containing a 1% of a solution of 10,000 units/mL penicillin and 10,000 mg/mL streptomycin). At 12 hours post infection, cells were washed once with PBS, and non-integrated viral cDNA was isolated from cells using a miniprep. Each mutant virus library was also subjected to a mock selection (no serum), and duplicate four 10-fold serial dilutions of each mutant virus library were also infected into 1×10^6^ cells to serve as an infectivity standard curve (Figure 1-Figure supplement 1). The proportion of the library that survived neutralization and entered cells was quantified using a qPCR and interpolation of the infectivity standard curve (Dingens et al., 2019). Sequencing libraries were generated using a barcoded subamplicon sequencing approach as previously described (Haddox et al., 2018, 2016) and detailed at https://jbloomlab.github.io/dms_tools2/bcsubamp.html. Libraries were sequenced using 2×250bp paired-end Illumina HiSeq runs.

Data was analyzed with dms_tools2 version 2.2.6 (https://jbloomlab.github.io/dms_tools2/) (Bloom, 2015). Differential selection is the log-transformed enrichment of a mutation relative to wildtype in the sera-selected condition relative to the non-selected control condition, corrected for sequencing depth and error. See prior work (Dingens et al., 2017; Doud et al., 2017) or https://jbloomlab.github.io/dms_tools2/diffsel.html for additional details. Sequencing of wildtype plasmid served DNA served as the error control while calculating differential selection statistics.

STR profiling on our stock of SupT1.CCR5 cells found that 11 of 14 alleles plus both amelogenin alleles matched the reference of the parental SupT1 cells (ATCC #CRL-1942) reference profile. These cells also tested negative for mycoplasma.

### EMPEM

IgG from rabbit sera was affinity-purified with equal parts of Protein G Sepharose (Sigma-Aldrich, P3296) and 150 ml Protein A Sepharose (GE, 17-5138-01) in a Poly-Prep Chromatography Column (BioRad, 731-1550). Fabs were generated by Papain cleavage and purified by the use of the Pierce Fab Preparation Kit according to the manufacturers’ instructions (Thermo Scientific - #44985). The purity of the Fabs was confirmed by SDS-PAGE.

BG505 SOSIP.664 or BG505 SOSIP v4.1/Fab complexes were made by mixing 15 μg SOSIP with 1 mg of polyclonal Fabs and allowed to incubate for 18 to 24 hours at room temperature. Complex samples they were SEC purified using a Superose 6 Increase 10/300 GL (GE Healthcare) column to remove excess Fab prior to EM grid preparation. Fractions containing the SOSIP/Fab complexes were pooled and concentrated using 10 kDa Amicon spin concentrators (Millipore). Samples were diluted to 0.02 mg/mL in TBS (0.05 M Tris pH 7.4, 0.15 M NaCl) and adsorbed onto glow discharged carbon-coated Cu400 EM grids (Electron Microscopy Sciences) and blotted after 10 seconds. The grids were then stained with 3 μL of 2% (w/v) uranyl formate, immediately blotted, waiting for 10 seconds before being stained again for 35 seconds followed by a final blot. Image collection and data processing was performed on TFS Talos F200C microscope (1.98 Å/pixel; 73,000× magnification) with an electron dose of ∼25 electrons/Å2 using Leginon (Pugach et al., 2015; Suloway et al., 2005). 2D classification, 3D sorting and 3D refinement was conducted using Relion v3.0 (Zivanov et al., 2018). EM density maps were visualized using UCSF Chimera (Pettersen et al., 2004) and segmented using Segger (Pintilie et al., 2010). Figures were generated using UCSF Chimera (Pettersen et al., 2004).

### TZM-bl neutralization assay

TZM-bl neutralization assays used to preliminarily map the specificity of our sera panel (Figure 1A) were performed in laboratories at Weill Cornell as previously described (Klasse et al., 2018). The remainder of the TZM-bl neutralization assays were performed in laboratories at the Fred Hutch as previously described (Dingens et al., 2017). These protocols are very similar and based on widely-used protocols TZM-bl neutralization assay protocols (Sarzotti-Kelsoe et al., 2014). Validation assays were completed in technical duplicate two to four times. In a small number of instances, the top of the neutralization plateau did not fit for single replicates; while fold change in ID_50_ values were always calculated using at minimum 2 replicates (with standard deviations (SD) between biological replicates reported), the MNP with SD reported as “(*)” were from single replicates in which the naturalization plateau was fit accurately.

## Data Availability

The entire mutational antigenic profiling analysis pipeline, as well as processed data are available as https://github.com/jbloomlab/Vacc_Rabbit_Sera_MAP. Illumina sequencing read were uploaded to the NCBI SRA as BioProject PRJNA656582 with sample identifiers SRR12431153-SRR12431189. EMPEM 3D maps are being processed to be deposited into EMDB.

## Acknowledgements

We thank Caelan Radford for comments on this manuscript and the Fred Hutch Genomics Core for performing the Illumina sequencing. We thank Gargi Debnath and Erik Francomano for assistance purifying and digesting IgG from sera.

## Author contributions

A.S.D.: Conceptualization, Investigation, Formal Analysis, Software, Writing—original draft, Writing—review & editing. P.P.: Investigation, Formal Analysis, Software, Writing—review & editing. K.D.M. – Investigation. S.K.H.: Software, Visualization, Data curation. T.K.: Investigation, Resources. C.C.: Resources. J.O.: Conceptualization, Writing—review & editing. J.P.M.: Conceptualization, Resources, Writing—review & editing. P.J.K.: Conceptualization, Resources, Writing—review & editing. A.B.W.: Conceptualization, Supervision, Funding, Writing—review & editing. J.D.B.: Conceptualization, Software, Supervision, Funding, Writing—review & editing.

## Supplemental Items

**Figure 1-Figure supplement 1.**
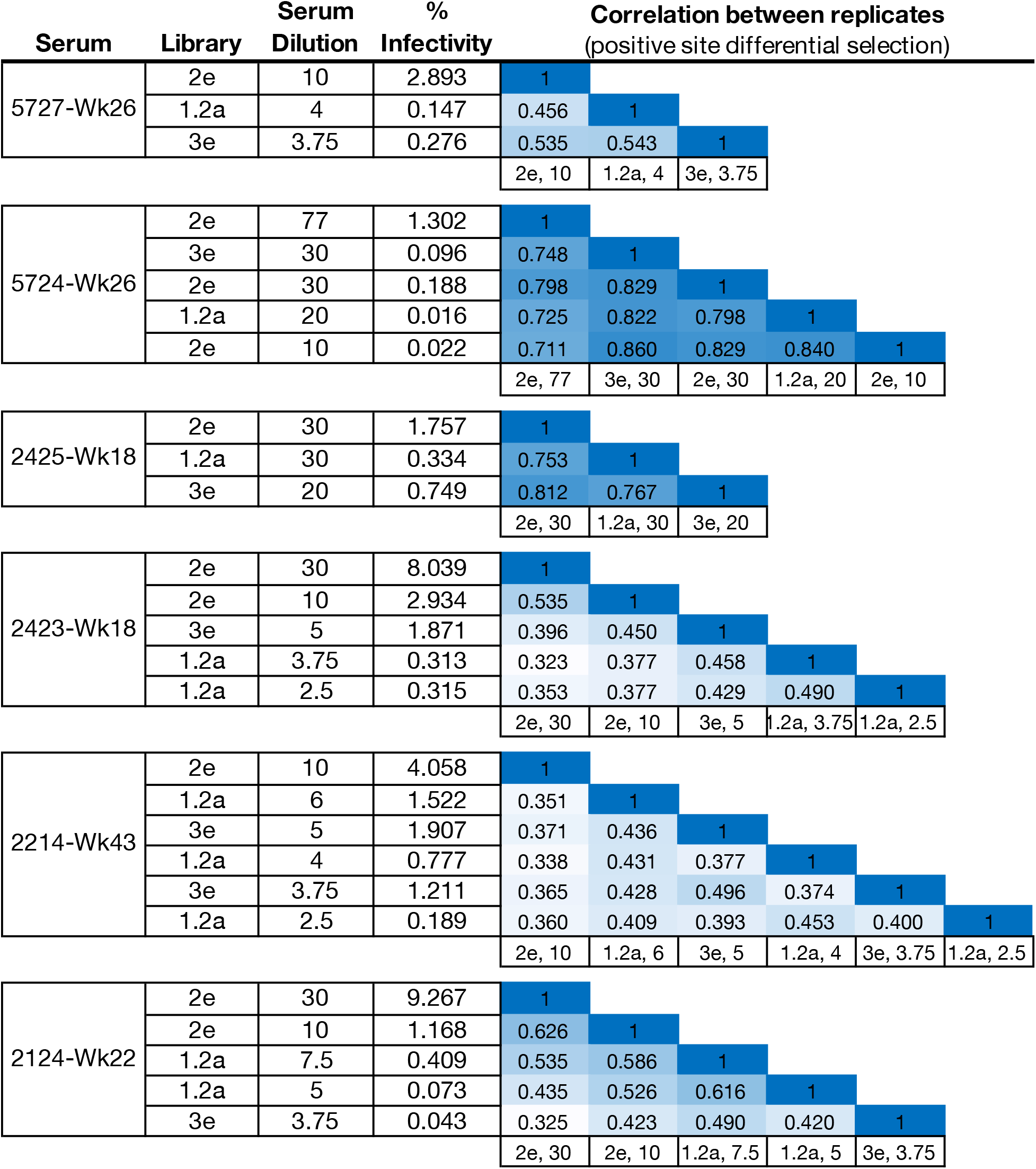
Experimental details for all mutational antigenic profiling experiments. The Library is the stock of independently passaged mutant virus library used in each experiment. The percentage of the mutant virus library that successfully entered cells after serum selection (% Infectivity) was measured using qPCR standard curves. The correlation matrix shows the Pearson’s r correlation coefficient of the positive site differential selection values between replicates.

**Figure 2–Figure supplement 1-6**. Mutational antigenic profiling data plotted for each individual replicates of pre- and post-immunization sera, grouped by rabbit. Data are plotted as in Figures 2A and 2B for each, but with mutations colored according to their biochemical property rather than epitope. (provided as a zip file in this single-PDF version)

**Figure 2-figure supplement 7.**
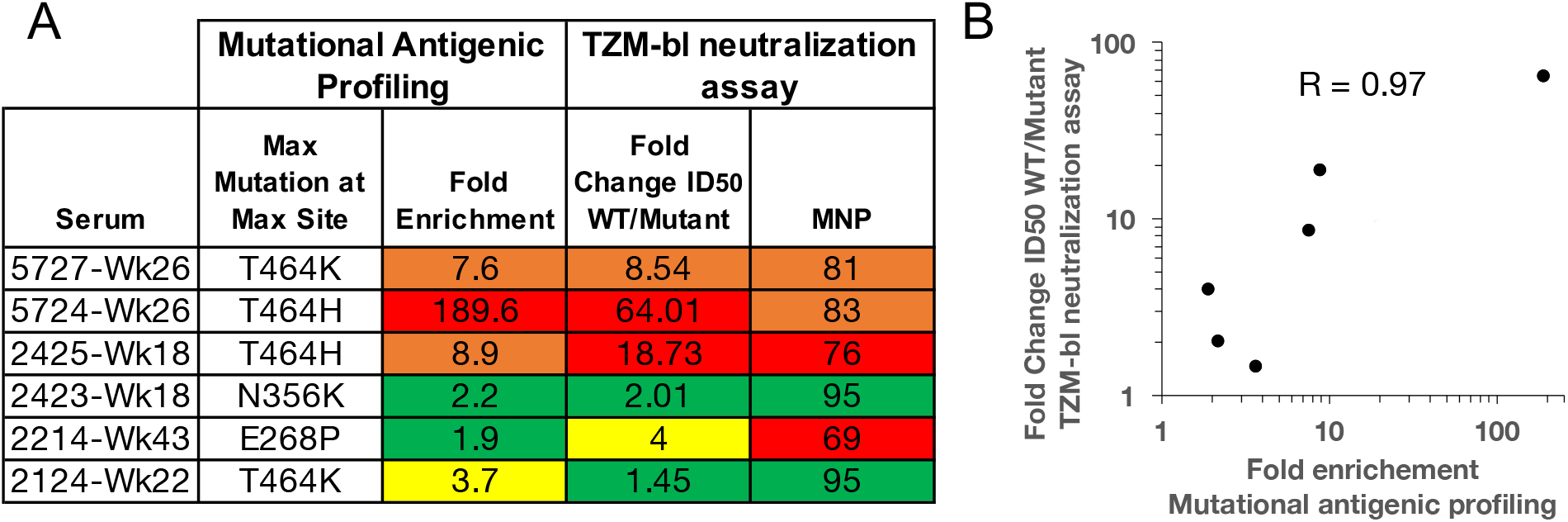
Validation of mutational antigenic profiling in neutralization assays. **A.** The fold enrichment in mutational antigenic profiling and the fold change in ID_50_ relative to wildtype and maximum neutralization plateau (MNP) from TZM-bl neutralization assays for the maximal effect mutant at the maximal effect site. The color scheme is as in Figure 1A **B.** Correlation between fold change ID_50_ relative to wildtype in TZM-bl neutralization assays and the fold enrichment in mutational antigenic profiling. The Pearson’s correlation coefficient is shown.

**Figure 2–Source Data 1.** Zip file containing csv files with all median site- and mutation-level differential selection values, as well as logoplots plotting escape profiles for the entire mutagenized portion of *env*.

**Figure 3-Figure supplement 1.**
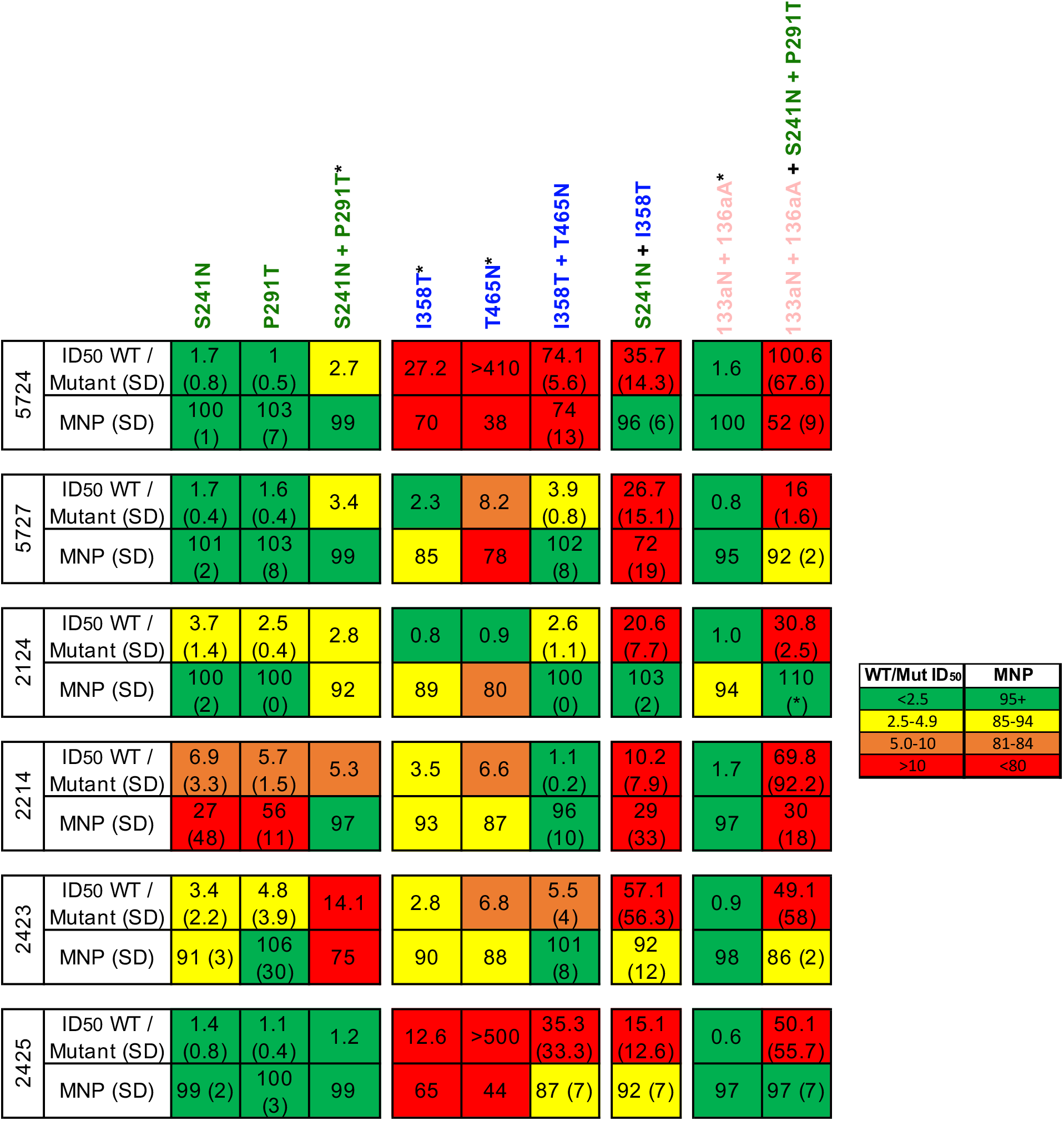
Effect of mutations that disrupt the C3/V5, glycan hole, and V1 epitopes alone and in combination with other epitope knockouts. Mutations are colored according to epitope, as in Figure 3. The standard deviation of replicate measures is shown in parentheses. Mutants labeled with an (*) are from the preliminary point mutant mapping in Figure 1A; these experiments were performed in a different lab than the remainder of TZM-bl point mutant mapping. While fold change in ID_50_ was always compared to a wildtype virus ran in parallel in independent labs, results across labs should only be interpreted generally (as presented in Figure 3C).

**Figure 4-Figure supplement 1.**
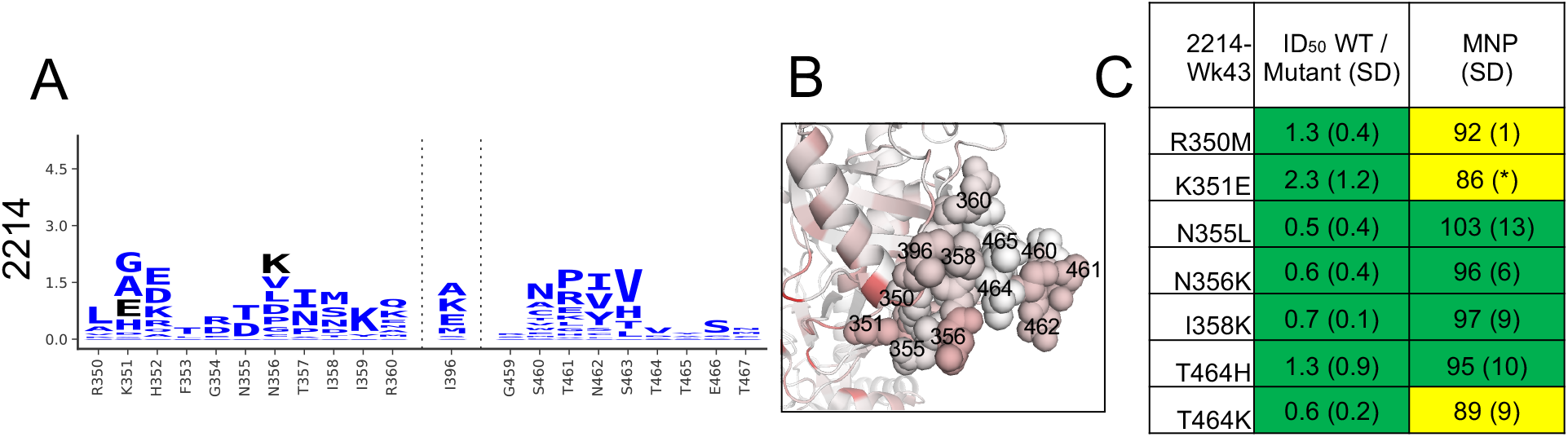
A, B, and C as in Figure 4, but for the single sera with limited targeting of the C3/V5 epitope.

**Figure 5-Figure supplement 1.**
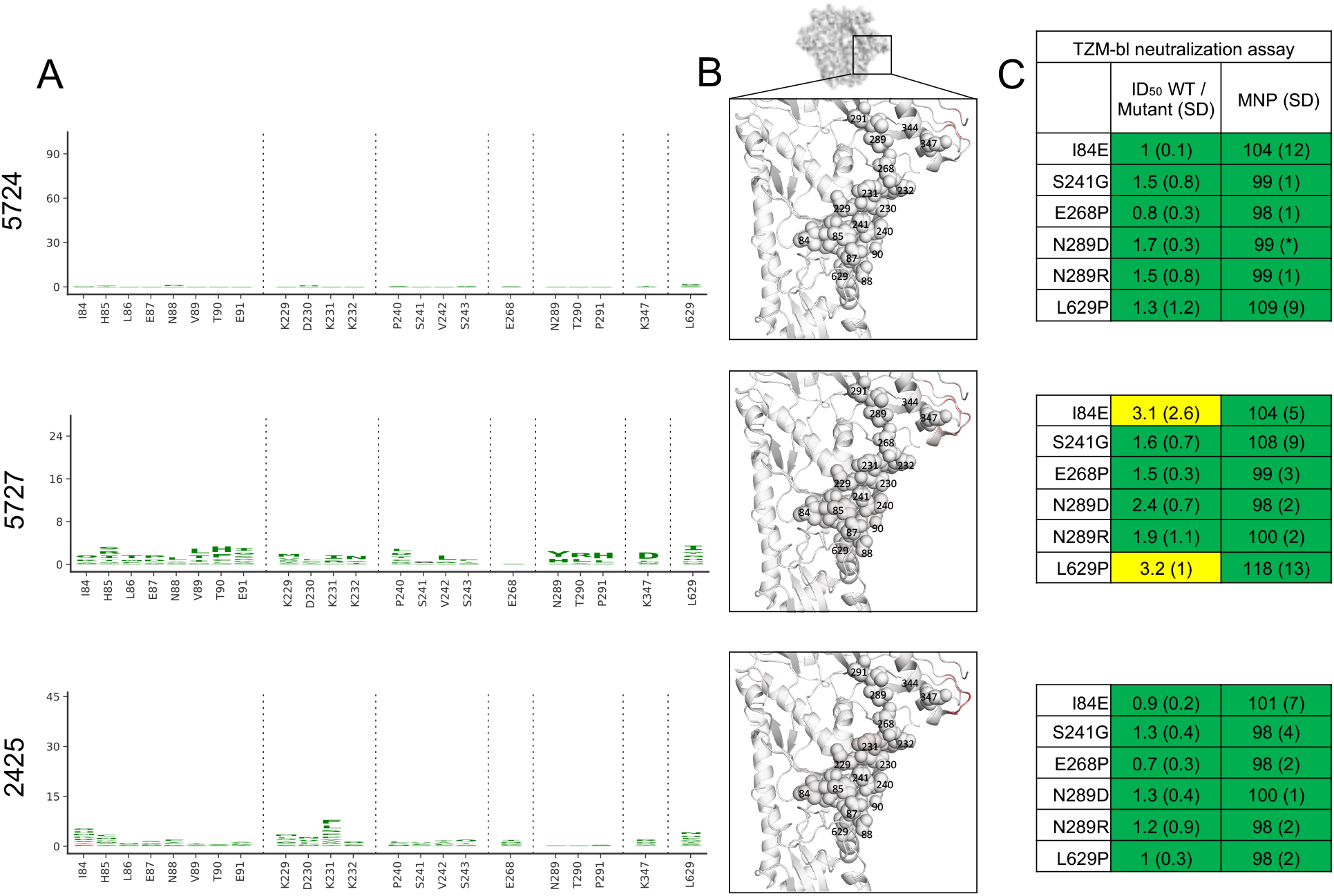
A, B, and C as in Figure 5, but for the three sera with limited targeting of the glycan hole epitope.

## References

Ackerman ME, Barouch DH, Alter G. 2017. Systems serology for evaluation of HIV vaccine trials. Immunol Rev 275:262–270. doi:10.1111/imr.12503

Alam SM, Aussedat B, Vohra Y, Meyerhoff RR, Cale EM, Walkowicz WE, Radakovich NA, Anasti K, Armand L, Parks R, Sutherland L, Scearce R, Joyce MG, Pancera M, Druz A, Georgiev IS, Von Holle T, Eaton A, Fox C, Reed SG, Louder M, Bailer RT, Morris L, Abdool-Karim SS, Cohen M, Liao HX, Montefiori DC, Park PK, Fernández-Tejada A, Wiehe K, Santra S, Kepler TB, Saunders KO, Sodroski J, Kwong PD, Mascola JR, Bonsignori M, Moody MA, Danishefsky S, Haynes BF. 2017. Mimicry of an HIV broadly neutralizing antibody epitope with a synthetic glycopeptide. Sci Transl Med. doi:10.1126/scitranslmed.aai7521

Barnes CO, West AP, Huey-Tubman KE, Hoffmann MAG, Sharaf NG, Hoffman PR, Koranda N, Gristick HB, Gaebler C, Muecksch F, Lorenzi JCC, Finkin S, Hägglöf T, Hurley A, Millard KG, Weisblum Y, Schmidt F, Hatziioannou T, Bieniasz PD, Caskey M, Robbiani DF, Nussenzweig MC, Bjorkman PJ. 2020. Structures of Human Antibodies Bound to SARS-CoV-2 Spike Reveal Common Epitopes and Recurrent Features of Antibodies. Cell. doi:10.1016/j.cell.2020.06.025

Bianchi M, Turner HL, Nogal B, Burton DR, Ward AB, Hangartner L, Bianchi M, Turner HL, Nogal B, Cottrell CA, Oyen D, Pauthner M. 2018. Electron-Microscopy-Based Epitope Mapping Defines Specificities of Polyclonal Antibodies Elicited during HIV-1 BG505 Envelope Trimer Article Electron-Microscopy-Based Epitope Mapping Defines Specificities of Polyclonal Antibodies Elicited during HIV-1 BG5. Immunity 49:288–300.e8. doi:10.1016/j.immuni.2018.07.009

Bloom JD. 2015. Software for the analysis and visualization of deep mutational scanning data. BMC Bioinformatics 16:1–13. doi:10.1186/s12859-015-0590-4

Boyoglu-Barnum S, Ellis D, Gillespie RA, Hutchinson GB, Park Y-J, Moin SM, Acton O, Ravichandran R, Murphy M, Pettie D, Matheson N, Carter L, Creanga A, Watson MJ, Kephart S, Vaile JR, Ueda G, Crank MC, Stewart L, Lee KK, Guttman M, Baker D, Mascola JR, Veesler D, Graham B, King NP, Kanekiyo M. 2020. Elicitation of broadly protective immunity to influenza by multivalent hemagglutinin nanoparticle vaccines. bioRxiv.

Briney B, Sok D, Jardine JG, Kulp DW, Skog P, Menis S, Jacak R, Kalyuzhniy O, de Val N, Sesterhenn F, Le KM, Ramos A, Jones M, Saye-Francisco KL, Blane TR, Spencer S, Georgeson E, Hu X, Ozorowski G, Adachi Y, Kubitz M, Sarkar A, Wilson IA, Ward AB, Nemazee D, Burton DR, Schief WR. 2016. Tailored Immunogens Direct Affinity Maturation toward HIV Neutralizing Antibodies. Cell 166:1459–1470.e11. doi:10.1016/j.cell.2016.08.005

Burton DR, Williamson RA, Parren PWHI. 2000. Antibody and virus: Binding and neutralization. Virology 270:1–3. doi:10.1006/viro.2000.0239

Chung AW, Kumar MP, Arnold KB, Yu WH, Schoen MK, Dunphy LJ, Suscovich TJ, Frahm N, Linde C, Mahan AE, Hoffner M, Streeck H, Ackerman ME, McElrath MJ, Schuitemaker H, Pau MG, Baden LR, Kim JH, Michael NL, Barouch DH, Lauffenburger D a., Alter G. 2015. Dissecting Polyclonal Vaccine-Induced Humoral Immunity against HIV Using Systems Serology. Cell 163:988–998. doi:10.1016/j.cell.2015.10.027

Correia BE, Bates JT, Loomis RJ, Baneyx G, Carrico C, Jardine JG, Rupert P, Correnti C, Kalyuzhniy O, Vittal V, Connell MJ, Stevens E, Schroeter A, Chen M, Macpherson S, Serra AM, Adachi Y, Holmes M a, Li Y, Klevit RE, Graham BS, Wyatt RT, Baker D, Strong RK, Crowe JE, Johnson PR, Schief WR. 2014. Proof of principle for epitope-focused vaccine design. Nature 507:201–6. doi:10.1038/nature12966

Cottrell CA, Schooten J van, Bowman CA, Yuan M, Oyen D, Shin M, Morpurgo R, Woude P van der, Breemen M van, Torres JL, Patel R, Gross J, Sewall LM, Copps J, Ozorowski G, Nogal B, Sok D, Rakasz EG, Labranche C, Vigdorovich V, Christley S, Carnathan DG, Sather DN, Montefiori D, Silvestri G, Burton DR, John P. Moore I, Wilson an A, Sanders RW, Ward AB, Gils MJ van. 2020. Mapping the immunogenic landscape of near-native HIV-1 envelope trimers in non-human primates. bioRxiv. doi:10.1101/2020.02.05.936096

De Taeye SW, Ozorowski G, Torrents De La Peña A, Guttman M, Julien JP, Van Den Kerkhof TLGM, Burger J a., Pritchard LK, Pugach P, Yasmeen A, Crampton J, Hu J, Bontjer I, Torres JL, Arendt H, Destefano J, Koff WC, Schuitemaker H, Eggink D, Berkhout B, Dean H, Labranche C, Crotty S, Crispin M, Montefiori DC, Klasse PJ, Lee KK, Moore JP, Wilson I a., Ward AB, Sanders RW. 2015. Immunogenicity of Stabilized HIV-1 Envelope Trimers with Reduced Exposure of Non-neutralizing Epitopes. Cell 163:1702–1715. doi:10.1016/j.cell.2015.11.056

Derking R, Allen JD, Cottrell CA, Sliepen K, Seabright GE, Lee W-H, Rantalainen K, Antanasijevic A, Copps J, Yasmeen A, Woude P van der, Taeye SW de, Kerkhof TLGM van den, Klasse PJ, Ozorowski G, Gils MJ van, Moore JP, Ward AB, Crispin M, Sanders RW. 2020. Enhancing glycan occupancy of soluble HIV-1 envelope trimers to mimic the native viral spike. bioRxiv. doi:10.1101/2020.07.02.184135

Dingens AS, Arenz D, Weight H, Overbaugh J, Bloom JD. 2019. An Antigenic Atlas of HIV-1 Escape from Broadly Neutralizing Antibodies Distinguishes Functional and Structural Epitopes. Immunity 50:520–532.e3. doi:10.1016/j.immuni.2018.12.017

Dingens AS, Haddox HK, Overbaugh J, Bloom JD. 2017. Comprehensive Mapping of HIV-1 Escape from a Broadly Neutralizing Antibody. Cell Host Microbe 21:777–787.e4. doi:10.1016/j.chom.2017.05.003

Dosenovic P, Von Boehmer L, Escolano A, Jardine J, Freund NT, Gitlin AD, McGuire AT, Kulp DW, Oliveira T, Scharf L, Pietzsch J, Gray MD, Cupo A, Van Gils MJ, Yao KH, Liu C, Gazumyan A, Seaman MS, Björkman PJ, Sanders RW, Moore JP, Stamatatos L, Schief WR, Nussenzweig MC. 2015. Immunization for HIV-1 Broadly Neutralizing Antibodies in Human Ig Knockin Mice. Cell. doi:10.1016/j.cell.2015.06.003

Doud MB, Hensley SE, Bloom JD. 2017. Complete mapping of viral escape from neutralizing antibodies. PLOS Pathog 13:e1006271. doi:10.1371/journal.ppat.1006271

Dubrovskaya V, Tran K, Ozorowski G, Guenaga J, Wilson R, Bale S, Cottrell CA, Turner HL, Seabright G, O’Dell S, Torres JL, Yang L, Feng Y, Leaman DP, Vázquez Bernat N, Liban T, Louder M, McKee K, Bailer RT, Movsesyan A, Doria-Rose NA, Pancera M, Karlsson Hedestam GB, Zwick MB, Crispin M, Mascola JR, Ward AB, Wyatt RT. 2019. Vaccination with Glycan-Modified HIV NFL Envelope Trimer-Liposomes Elicits Broadly Neutralizing Antibodies to Multiple Sites of Vulnerability. Immunity 51:915–929.e7. doi:10.1016/j.immuni.2019.10.008

Escolano A, Gristick HB, Abernathy ME, Merkenschlager J, Gautam R, Oliveira TY, Pai J, West AP, Barnes CO, Cohen AA, Wang H, Golijanin J, Yost D, Keeffe JR, Wang Z, Zhao P, Yao KH, Bauer J, Nogueira L, Gao H, Voll A V., Montefiori DC, Seaman MS, Gazumyan A, Silva M, McGuire AT, Stamatatos L, Irvine DJ, Wells L, Martin MA, Bjorkman PJ, Nussenzweig MC. 2019. Immunization expands B cells specific to HIV-1 V3 glycan in mice and macaques. Nature 570:468–473. doi:10.1038/s41586-019-1250-z

Escolano A, Steichen JM, Dosenovic P, Kulp DW, Golijanin J, Sok D, Freund NT, Gitlin AD, Oliveira T, Araki T, Lowe S, Chen ST, Heinemann J, Yao K-H, Georgeson E, Saye-Francisco KL, Gazumyan A, Adachi Y, Kubitz M, Burton DR, Schief WR, Nussenzweig MC. 2016. Sequential Immunization Elicits Broadly Neutralizing Anti-HIV-1 Antibodies in Ig Knockin Mice. Cell 166:1445–1458.e12. doi:10.1016/j.cell.2016.07.030

Georgiev IS, Doria-Rose N a, Zhou T, Kwon Y Do, Staupe RP, Moquin S, Chuang G-Y, Louder MK, Schmidt SD, Altae-Tran HR, Bailer RT, McKee K, Nason M, O’Dell S, Ofek G, Pancera M, Srivatsan S, Shapiro L, Connors M, Migueles S a, Morris L, Nishimura Y, Martin M a, Mascola JR, Kwong PD. 2013. Delineating antibody recognition in polyclonal sera from patterns of HIV-1 isolate neutralization. Science 340:751–6. doi:10.1126/science.1233989

Haddox HK, Dingens AS, Bloom JD. 2016. Experimental Estimation of the Effects of All Amino-Acid Mutations to HIV’s Envelope Protein on Viral Replication in Cell Culture. PLOS Pathog 12:e1006114. doi:10.1371/journal.ppat.1006114

Haddox HK, Dingens AS, Hilton SK, Overbaugh J, Bloom JD. 2018. Mapping mutational effects along the evolutionary landscape of HIV envelope. Elife 7:e34420. doi:10.7554/eLife.34420

Haynes BF, Kelsoe G, Harrison SC, Kepler TB. 2012. B-cell-lineage immunogen design in vaccine development with HIV-1 as a case study. Nat Biotechnol. doi:10.1038/nbt.2197

Hilton SK, Huddleston J, Black A, North K, Dingens AS, Bedford T, Bloom JD. 2020. dms-view: Interactive visualization tool for deep mutational scanning data. bioRxiv 1–6. doi:10.1101/2020.05.14.096842

Hu JK, Crampton JC, Cupo A, Ketas T, van Gils MJ, Sliepen K, de Taeye SW, Sok D, Ozorowski G, Deresa I, Stanfield R, Ward AB, Burton DR, Klasse PJ, Sanders RW, Moore JP, Crotty S. 2015. Murine Antibody Responses to Cleaved Soluble HIV-1 Envelope Trimers Are Highly Restricted in Specificity. J Virol 89:10383–10398. doi:10.1128/jvi.01653-15

Jardine J, Julien J-P, Menis S, Ota T, Kalyuzhniy O, McGuire A, Sok D, Huang P-S, MacPherson S, Jones M, Nieusma T, Mathison J, Baker D, Ward AB, Burton DR, Stamatatos L, Nemazee D, Wilson IA, Schief WR. 2013. Rational HIV Immunogen Design to Target Specific Germline B Cell Receptors. Science (80-) 340:711–716. doi:10.1126/science.1234150

Jardine JG, Kulp DW, Havenar-Daughton C, Sarkar A, Briney B, Sok D, Sesterhenn F, Ereno-Orbea J, Kalyuzhniy O, Deresa I, Hu X, Spencer S, Jones M, Georgeson E, Adachi Y, Kubitz M, DeCamp AC, Julien J-P, Wilson IA, Burton DR, Crotty S, Schief WR. 2016. HIV-1 broadly neutralizing antibody precursor B cells revealed by germline-targeting immunogen. Science (80-) 351:1458–1463. doi:10.1126/science.aad9195

Julien JP, Cupo A, Sok D, Stanfield RL, Lyumkis D, Deller MC, Klasse PJ, Burton DR, Sanders RW, Moore JP, Ward AB, Wilson IA. 2013. Crystal structure of a soluble cleaved HIV-1 envelope trimer. Science (80-). doi:10.1126/science.1245625

Klasse PJ, Ketas TJ, Cottrell CA, Ozorowski G, Debnath G, Camara D, Francomano E, Pugach P, Ringe RP, LaBranche CC, van Gils MJ, Bricault CA, Barouch DH, Crotty S, Silvestri G, Kasturi S, Pulendran B, Wilson IA, Montefiori DC, Sanders RW, Ward AB, Moore JP. 2018. Epitopes for neutralizing antibodies induced by HIV-1 envelope glycoprotein BG505 SOSIP trimers in rabbits and macaques. PLoS Pathog 14:1–20. doi:10.1371/journal.ppat.1006913

Klasse PJ, LaBranche CC, Ketas TJ, Ozorowski G, Cupo A, Pugach P, Ringe RP, Golabek M, van Gils MJ, Guttman M, Lee KK, Wilson IA, Butera ST, Ward AB, Montefiori DC, Sanders RW, Moore JP. 2016. Sequential and Simultaneous Immunization of Rabbits with HIV-1 Envelope Glycoprotein SOSIP.664 Trimers from Clades A, B and C. PLoS Pathog 12:1–31. doi:10.1371/journal.ppat.1005864

Kong R, Duan H, Sheng Z, Xu K, Acharya P, Chen X, Cheng C, Dingens AS, Gorman J, Sastry M, Shen CH, Zhang B, Zhou T, Chuang GY, Chao CW, Gu Y, Jafari AJ, Louder MK, O’Dell S, Rowshan AP, Viox EG, Wang Y, Choi CW, Corcoran MM, Corrigan AR, Dandey VP, Eng ET, Geng H, Foulds KE, Guo Y, Kwon YD, Lin B, Liu K, Mason RD, Nason MC, Ohr TY, Ou L, Rawi R, Sarfo EK, Schön A, Todd JP, Wang S, Wei H, Wu W, Mullikin JC, Bailer RT, Doria-Rose NA, Karlsson Hedestam GB, Scorpio DG, Overbaugh J, Bloom JD, Carragher B, Potter CS, Shapiro L, Kwong PD, Mascola JR. 2019. Antibody Lineages with Vaccine-Induced Antigen-Binding Hotspots Develop Broad HIV Neutralization. Cell 567–584. doi:10.1016/j.cell.2019.06.030

Kreer C, Gruell H, Mora T, Walczak AM, Klein F. 2020. Exploiting B cell receptor analyses to inform on HIV-1 vaccination strategies. Vaccines 8:1–19. doi:10.3390/vaccines8010013

Kulp DW, Steichen JM, Pauthner M, Hu X, Schiffner T, Liguori A, Cottrell CA, Havenar-Daughton C, Ozorowski G, Georgeson E, Kalyuzhniy O, Willis JR, Kubitz M, Adachi Y, Reiss SM, Shin M, de Val N, Ward AB, Crotty S, Burton DR, Schief WR. 2017. Structure-based design of native-like HIV-1 envelope trimers to silence non-neutralizing epitopes and eliminate CD4 binding. Nat Commun 8:1655. doi:10.1038/s41467-017-01549-6

Kwong PD, Mascola JR. 2018. HIV-1 Vaccines Based on Antibody Identification, B Cell Ontogeny, and Epitope Structure. Immunity 48:855–871. doi:10.1016/j.immuni.2018.04.029

Lavinder JJ, Wine Y, Giesecke C, Ippolito GC, Horton AP, Lungu OI, Hoi KH, DeKosky BJ, Murrin EM, Wirth MM, Ellington AD, Dörner T, Marcotte EM, Boutz DR, Georgiou G. 2014. Identification and characterization of the constituent human serum antibodies elicited by vaccination. Proc Natl Acad Sci. doi:10.1073/pnas.1317793111

Lei L, Yang YR, Tran K, Wang Y, Chiang C-II, Ozorowski G, Xiao Y, Ward AB, Wyatt RT, Li Y. 2019. The HIV-1 Envelope Glycoprotein C3/V4 Region Defines a Prevalent Neutralization Epitope following Immunization. Cell Rep 27:586–598.e6. doi:10.1016/j.celrep.2019.03.039

Lyumkis D, Julien J-P, de Val N, Cupo A, Potter CS, Klasse P-J, Burton DR, Sanders RW, Moore JP, Carragher B, Wilson IA, Ward AB. 2013. Cryo-EM Structure of a Fully Glycosylated Soluble Cleaved HIV-1 Envelope Trimer. Science (80-) 342:1484–1490. doi:10.1126/science.1245627

McCoy LE, Van Gils MJ, Ozorowski G, Ward AB, Sanders RW, Burton DR, Messmer T, Briney B, Voss JE, Kulp DW, Macauley MS, Sok D, Pauthner M, Menis S, Cottrell C a, Torres JL, Hsueh J, Schief WR, Wilson I a. 2016. Holes in the Glycan Shield of the Native HIV Envelope Are a Target of Trimer-Elicited Neutralizing Antibodies. Cell Rep 16:1–12. doi:10.1016/j.celrep.2016.07.074

Medina-Ramírez M, Garces F, Escolano A, Skog P, de Taeye SW, Del Moral-Sanchez I, McGuire AT, Yasmeen A, Behrens A-J, Ozorowski G, van den Kerkhof TLGM, Freund NT, Dosenovic P, Hua Y, Gitlin AD, Cupo A, van der Woude P, Golabek M, Sliepen K, Blane T, Kootstra N, van Breemen MJ, Pritchard LK, Stanfield RL, Crispin M, Ward AB, Stamatatos L, Klasse PJ, Moore JP, Nemazee D, Nussenzweig MC, Wilson IA, Sanders RW. 2017. Design and crystal structure of a native-like HIV-1 envelope trimer that engages multiple broadly neutralizing antibody precursors in vivo. J Exp Med 214:2573–2590. doi:10.1084/jem.20161160

Nogal B, Bianchi M, Cottrell CA, Kirchdoerfer RN, Sewall LM, Turner HL, Zhao F, Sok D, Burton DR, Hangartner L, Ward AB. 2020. Mapping Polyclonal Antibody Responses in Non-human Primates Vaccinated with HIV Env Trimer Subunit Vaccines. Cell Rep 30:3755–3765.e7. doi:10.1016/j.celrep.2020.02.061

Nogal B, Mccoy LE, Gils MJ Van, Cottrell CA, James E, Andrabi R, Pauthner M, Liang C. 2019. HIV Envelope Trimer-Elicited Autologous Neutralizing Antibodies Bind a Region Overlapping the N332 Glycan Supersite. bioRxiv. doi:10.1101/831008

Pancera M, Zhou T, Druz A, Georgiev IS, Soto C, Gorman J, Huang J, Acharya P, Chuang G-Y, Ofek G, Stewart-Jones GBE, Stuckey J, Bailer RT, Joyce MG, Louder MK, Tumba N, Yang Y, Zhang B, Cohen MS, Haynes BF, Mascola JR, Morris L, Munro JB, Blanchard SC, Mothes W, Connors M, Kwong PD. 2014. Structure and immune recognition of trimeric pre-fusion HIV-1 Env. Nature 514:455–461. doi:10.1038/nature13808

Pauthner M, Sharma SK, Butera ST, Wilson IA, Sanders RW, Kaushik K, Havenar-Daughton C, Wyatt RT, Montefiori DC, Eroshkin AM, Post KW, Steichen JM, Kulp DW, Oom AL, Chandrashekar A, Silvestri G, Tokatlian T, Burton DR, Schief WR, Barouch DH, Cottrell CA, Ward AB, Irvine DJ, de Taeye SW, Carnathan DG, Nkolola JP, LaBranche CC, Boopathy A V., Bastidas R, Liu J, McCoy LE, Crotty S, Ozorowski G, Guenaga J, Torrents de la Peña A, Sok D, Cirelli KM. 2017. Elicitation of Robust Tier 2 Neutralizing Antibody Responses in Nonhuman Primates by HIV Envelope Trimer Immunization Using Optimized Approaches. Immunity 46:1073–1088.e6. doi:10.1016/j.immuni.2017.05.007

Pauthner MG, Nkolola JP, Havenar-Daughton C, Murrell B, Reiss SM, Bastidas R, Prévost J, Nedellec R, von Bredow B, Abbink P, Cottrell CA, Kulp DW, Tokatlian T, Nogal B, Bianchi M, Li H, Lee JH, Butera ST, Evans DT, Hangartner L, Finzi A, Wilson IA, Wyatt RT, Irvine DJ, Schief WR, Ward AB, Sanders RW, Crotty S, Shaw GM, Barouch DH, Burton DR. 2019. Vaccine-Induced Protection from Homologous Tier 2 SHIV Challenge in Nonhuman Primates Depends on Serum-Neutralizing Antibody Titers. Immunity 50:241-252.e6. doi:10.1016/j.immuni.2018.11.011

Pettersen EF, Goddard TD, Huang CC, Couch GS, Greenblatt DM, Meng EC, Ferrin TE. 2004. UCSF Chimera—A Visualization System for Exploratory Research and Analysis. J Comput Chem 25:1605–1612. doi:10.1002/jcc.20084

Pintilie GD, Zhang J, Goddard TD, Chiu W, Gossard DC. 2010. Quantitative analysis of cryo-EM density map segmentation by watershed and scale-space filtering, and fitting of structures by alignment to regions. J Struct Biol. doi:10.1016/j.jsb.2010.03.007

Pugach P, Ozorowski G, Cupo A, Ringe R, Yasmeen A, de Val N, Derking R, Kim HJ, Korzun J, Golabek M, de los Reyes K, Ketas TJ, Julien J-P, Burton DR, Wilson I a., Sanders RW, Klasse PJ, Ward AB, Moore JP. 2015. A native-like SOSIP.664 trimer based on a HIV-1 subtype B env gene. J Virol 89:JVI.03473-14. doi:10.1128/JVI.03473-14

Ringe RP, Ozorowski G, Rantalainen K, Struwe WB, Matthews K, Torres JL, Yasmeen A, Cottrell CA, Ketas TJ, LaBranche CC, Montefiori DC, Cupo A, Crispin M, Wilson IA, Ward AB, Sanders RW, Klasse PJ, Moore JP. 2017. Reducing V3 Antigenicity and Immunogenicity on Soluble, Native-Like HIV-1 Env SOSIP Trimers. J Virol 91:1–17. doi:10.1128/JVI.00677-17

Ringe RP, Pugach P, Cottrell CA, LaBranche CC, Seabright GE, Ketas TJ, Ozorowski G, Kumar S, Schorcht A, van Gils MJ, Crispin M, Montefiori DC, Wilson IA, Ward AB, Sanders RW, Klasse PJ, Moore JP. 2019. Closing and Opening Holes in the Glycan Shield of HIV-1 Envelope Glycoprotein SOSIP Trimers Can Redirect the Neutralizing Antibody Response to the Newly Unmasked Epitopes. J Virol 93:1–26. doi:10.1128/JVI.01656-18

Sanders RW, Derking R, Cupo A, Julien JP, Yasmeen A, de Val N, Kim HJ, Blattner C, Torrents de la Pena A, Korzun J, Golabek M, de los Reyes K, Ketas TJ, van Gils MJ, King CR, Wilson I a., Ward AB, Klasse PJ, Moore JP. 2013. A Next-Generation Cleaved, Soluble HIV-1 Env Trimer, BG505 SOSIP.664 gp140, Expresses Multiple Epitopes for Broadly Neutralizing but Not Non-Neutralizing Antibodies. PLoS Pathog 9:e1003618. doi:10.1371/journal.ppat.1003618

Sanders RW, Moore JP. 2017. Native-like Env trimers as a platform for HIV-1 vaccine design. Immunol Rev 275:161–182. doi:10.1111/imr.12481

Sanders RW, van Gils MJ, Derking R, Sok D, Ketas TJ, Burger J a., Ozorowski G, Cupo a., Simonich C, Goo L, Arendt H, Kim HJ, Lee JH, Pugach P, Williams M, Debnath G, Moldt B, van Breemen MJ, Isik G, Medina-Ramirez M, Back JW, Koff WC, Julien J-P, Rakasz EG, Seaman MS, Guttman M, Lee KK, Klasse PJ, LaBranche C, Schief WR, Wilson I a., Overbaugh J, Burton DR, Ward a. B, Montefiori DC, Dean H, Moore JP. 2015. HIV-1 neutralizing antibodies induced by native-like envelope trimers. Science (80-) science.aac4223-. doi:10.1126/science.aac4223

Sarzotti-Kelsoe M, Bailer RT, Turk E, Lin CL, Bilska M, Greene KM, Gao H, Todd C a., Ozaki D a., Seaman MS, Mascola JR, Montefiori DC. 2014. Optimization and validation of the TZM-bl assay for standardized assessments of neutralizing antibodies against HIV-1. J Immunol Methods 409:131–146. doi:10.1016/j.jim.2013.11.022

Saunders KO, Verkoczy LK, Jiang C, Zhang J, Parks R, Chen H, Housman M, Bouton-Verville H, Shen X, Trama AM, Scearce R, Sutherland L, Santra S, Newman A, Eaton A, Xu K, Georgiev IS, Joyce MG, Tomaras GD, Bonsignori M, Reed SG, Salazar A, Mascola JR, Moody MA, Cain DW, Centlivre M, Zurawski S, Zurawski G, Erickson HP, Kwong PD, Alam SM, Levy Y, Montefiori DC, Haynes BF. 2017. Vaccine Induction of Heterologous Tier 2 HIV-1 Neutralizing Antibodies in Animal Models. Cell Rep. doi:10.1016/j.celrep.2017.12.028

Scheid JF, Mouquet H, Feldhahn N, Seaman MS, Velinzon K, Pietzsch J, Ott RG, Anthony RM, Zebroski H, Hurley A, Phogat A, Chakrabarti B, Li Y, Connors M, Pereyra F, Walker BD, Wardemann H, Ho D, Wyatt RT, Mascola JR, Ravetch J V, Nussenzweig MC. 2009. Broad diversity of neutralizing antibodies isolated from memory B cells in HIV-infected individuals. Nature 458:636–640. doi:10.1038/nature07930

Steichen J M, Steichen Jon M, Lin Y, Havenar-daughton C, Pecetta S, Ozorowski G, Willis JR, Toy L, Sok D, Liguori A, Kratochvil S, Torres JL, Kalyuzhniy O, Melzi E, Kulp DW, Raemisch S, Hu X, Bernard SM, Georgeson E, Phelps N, Adachi Y, Kubitz M, Landais E, Umotoy J, Robinson A, Briney B, Wilson IA, Burton DR, Ward AB, Crotty S, Batista FD, Schief WR. 2019. A generalized HIV vaccine design strategy for priming of broadly neutralizing antibody responses 4380:1–23.

Steichen JM, Kulp DW, Tokatlian T, Escolano A, Dosenovic P, Stanfield RL, McCoy LE, Ozorowski G, Hu X, Kalyuzhniy O, Briney B, Schiffner T, Garces F, Freund NT, Gitlin AD, Menis S, Georgeson E, Kubitz M, Adachi Y, Jones M, Mutafyan AA, Yun DS, Mayer CT, Ward AB, Burton DR, Wilson IA, Irvine DJ, Nussenzweig MC, Schief WR. 2016. HIV Vaccine Design to Target Germline Precursors of Glycan-Dependent Broadly Neutralizing Antibodies. Immunity 45:483–496. doi:10.1016/j.immuni.2016.08.016

Suloway C, Pulokas J, Fellmann D, Cheng A, Guerra F, Quispe J, Stagg S, Potter CS, Carragher B. 2005. Automated molecular microscopy: The new Leginon system. J Struct Biol. doi:10.1016/j.jsb.2005.03.010

Tian M, Cheng C, Chen X, Duan H, Cheng H-L, Dao M, Sheng Z, Kimble M, Wang L, Lin S, Schmidt SD, Du Z, Joyce MG, Chen Yiwei, DeKosky BJ, Chen Yimin, Normandin E, Cantor E, Chen RE, Doria-Rose NA, Zhang Y, Shi W, Kong W-P, Choe M, Henry AR, Laboune F, Georgiev IS, Huang P-Y, Jain S, McGuire AT, Georgeson E, Menis S, Douek DC, Schief WR, Stamatatos L, Kwong PD, Shapiro L, Haynes BF, Mascola JR, Alt FW. 2016. Induction of HIV Neutralizing Antibody Lineages in Mice with Diverse Precursor Repertoires. Cell 166:1471–1484.e18. doi:10.1016/j.cell.2016.07.029

Torrents de la Peña A, de Taeye SW, Sliepen K, LaBranche CC, Burger JA, Schermer EE, Montefiori DC, Moore JP, Klasse PJ, Sanders RW. 2018. Immunogenicity in Rabbits of HIV-1 SOSIP Trimers from Clades A, B, and C, Given Individually, Sequentially, or in Combination. J Virol. doi:10.1128/jvi.01957-17

Torrents de la Peña A, Julien JP, de Taeye SW, Garces F, Guttman M, Ozorowski G, Pritchard LK, Behrens AJ, Go EP, Burger JA, Schermer EE, Sliepen K, Ketas TJ, Pugach P, Yasmeen A, Cottrell CA, Torres JL, Vavourakis CD, van Gils MJ, LaBranche C, Montefiori DC, Desaire H, Crispin M, Klasse PJ, Lee KK, Moore JP, Ward AB, Wilson IA, Sanders RW. 2017. Improving the Immunogenicity of Native-like HIV-1 Envelope Trimers by Hyperstabilization. Cell Rep 20:1805–1817. doi:10.1016/j.celrep.2017.07.077

Ward AB, Wilson IA. 2020. Innovations in structure-based antigen design and immune monitoring for next generation vaccines. Curr Opin Immunol 116544. doi:10.1016/j.coi.2020.03.013

Wine Y, Boutz DR, Lavinder JJ, Miklos AE, Hughes R a, Hoi KH, Jung ST, Horton AP, Murrin EM, Ellington AD, Marcotte EM, Georgiou G. 2013. Molecular deconvolution of the monoclonal antibodies that comprise the polyclonal serum response. Proc Natl Acad Sci U S A 110:2993–8. doi:10.1073/pnas.1213737110

Wu X, Parast AB, Richardson BA, Nduati R, John-Stewart G, Mbori-Ngacha D, Rainwater SMJ, Overbaugh J. 2006. Neutralization Escape Variants of Human Immunodeficiency Virus Type 1 Are Transmitted from Mother to Infant 80:835–844. doi:10.1128/JVI.80.2.835

Xu K, Acharya P, Kong R, Cheng C, Chuang G-Y, Liu K, Louder MK, O’Dell S, Rawi R, Sastry M, Shen C-H, Zhang B, Zhou T, Asokan M, Bailer RT, Chambers M, Chen X, Choi CW, Dandey VP, Doria-Rose NA, Druz A, Eng ET, Farney SK, Foulds KE, Geng H, Georgiev IS, Gorman J, Hill KR, Jafari AJ, Kwon YD, Lai Y-T, Lemmin T, McKee K, Ohr TY, Ou L, Peng D, Rowshan AP, Sheng Z, Todd J-P, Tsybovsky Y, Viox EG, Wang Y, Wei H, Yang Y, Zhou AF, Chen R, Yang L, Scorpio DG, McDermott AB, Shapiro L, Carragher B, Potter CS, Mascola JR, Kwong PD, Kai Xu*, Acharya* P, Kong* R, Cheng* C, Chuang G-Y, Liu K, Louder MK, O’Dell S, Rawi R, Sastry M, Shen C-H, Zhang B, Zhou T, Asokan M, Bailer RT, Chambers M, Chen X, Choi CW, Dandey VP, Doria-Rose NA, Druz A, Eng ET, Farney K, Foulds KE, Geng H, Georgiev IS, Gorman J, Hill KR, Jafari AJ, Kwon YD, Lai Y-T, Lemmin T, McKee K, Ohr TY, Ou L, Peng D, P. A, Roshan, Sheng Z, Todd J-P, Tsybovsky Y, Viox EG, Wang Y, Wei H, Yang Y, Zhou AF, Chen R, Yang L, Scorpio DG, McDermott AB, Shapiro L, Carragher B, Potter CS, Mascola JR, Kwong PD, Xu K, Acharya P, Kong R, Cheng C. 2018. Epitope-based vaccine design yields fusion peptide-directed antibodies that neutralize diverse strains of HIV-1. Nat Med 24:857–867. doi:10.1038/s41591-018-0042-6

Yang X, Lipchina I, Cocklin S, Chaiken I, Sodroski J. 2006. Antibody Binding Is a Dominant Determinant of the Efficiency of Human Immunodeficiency Virus Type 1 Neutralization. J Virol 80:11404–11408. doi:10.1128/jvi.01102-06

Yang YR, McCoy LE, van Gils MJ, Andrabi R, Turner HL, Yuan M, Cottrell CA, Ozorowski G, Voss J, Pauthner M, Polveroni TM, Messmer T, Wilson IA, Sanders RW, Burton DR, Ward AB. 2020. Autologous Antibody Responses to an HIV Envelope Glycan Hole Are Not Easily Broadened in Rabbits. J Virol 94:1–15. doi:10.1128/jvi.01861-19

Zivanov J, Nakane T, Forsberg BO, Kimanius D, Hagen WJH, Lindahl E, Scheres SHW. 2018. New tools for automated high-resolution cryo-EM structure determination in RELION-3. Elife. doi:10.7554/eLife.42166

