## Supplementary figures and images for "High-resolution mapping of the neutralizing and binding specificities of polyclonal rabbit serum elicited by HIV Env trimer immunization"

### 2124.pdf

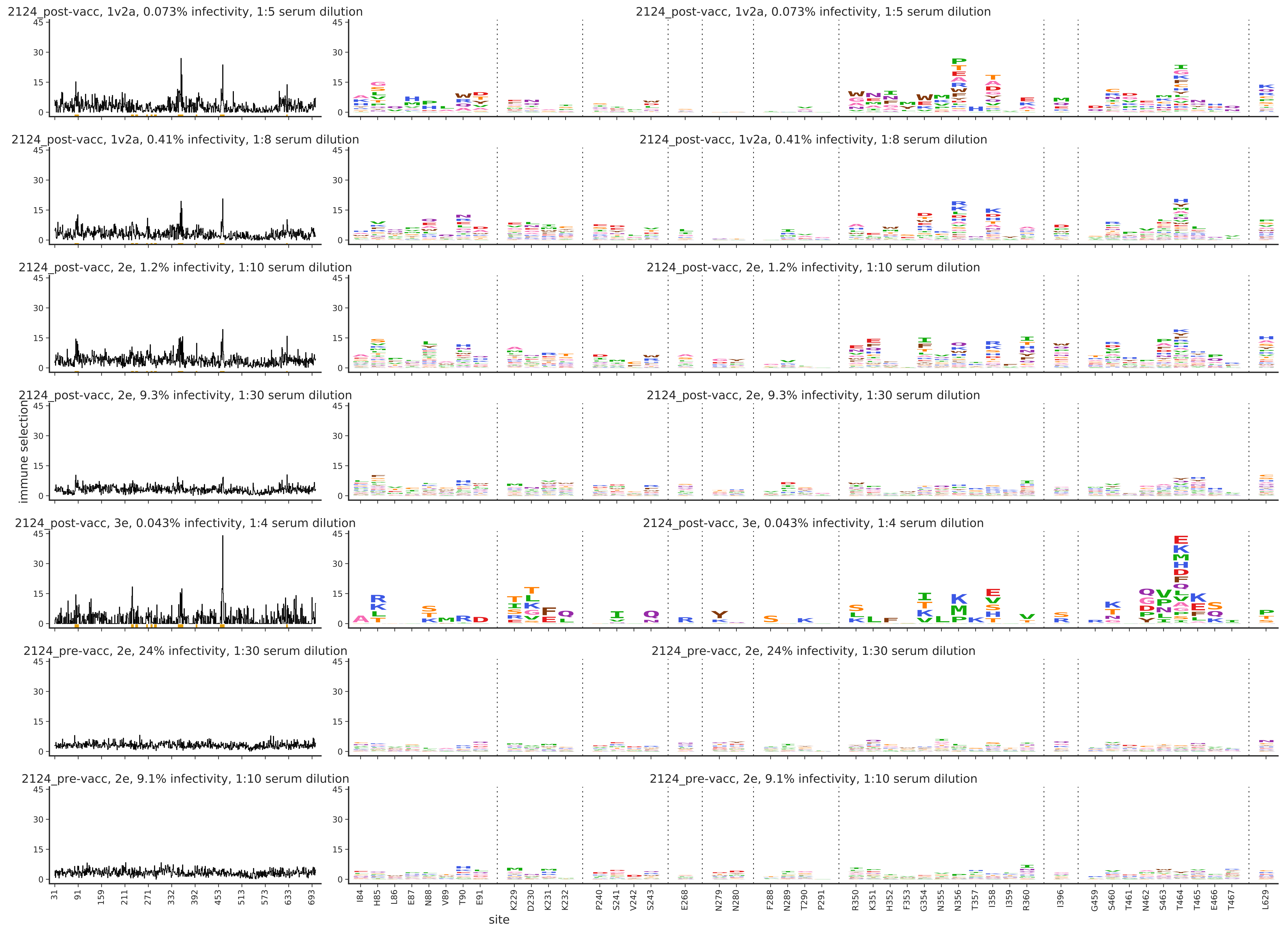

### 2214.pdf

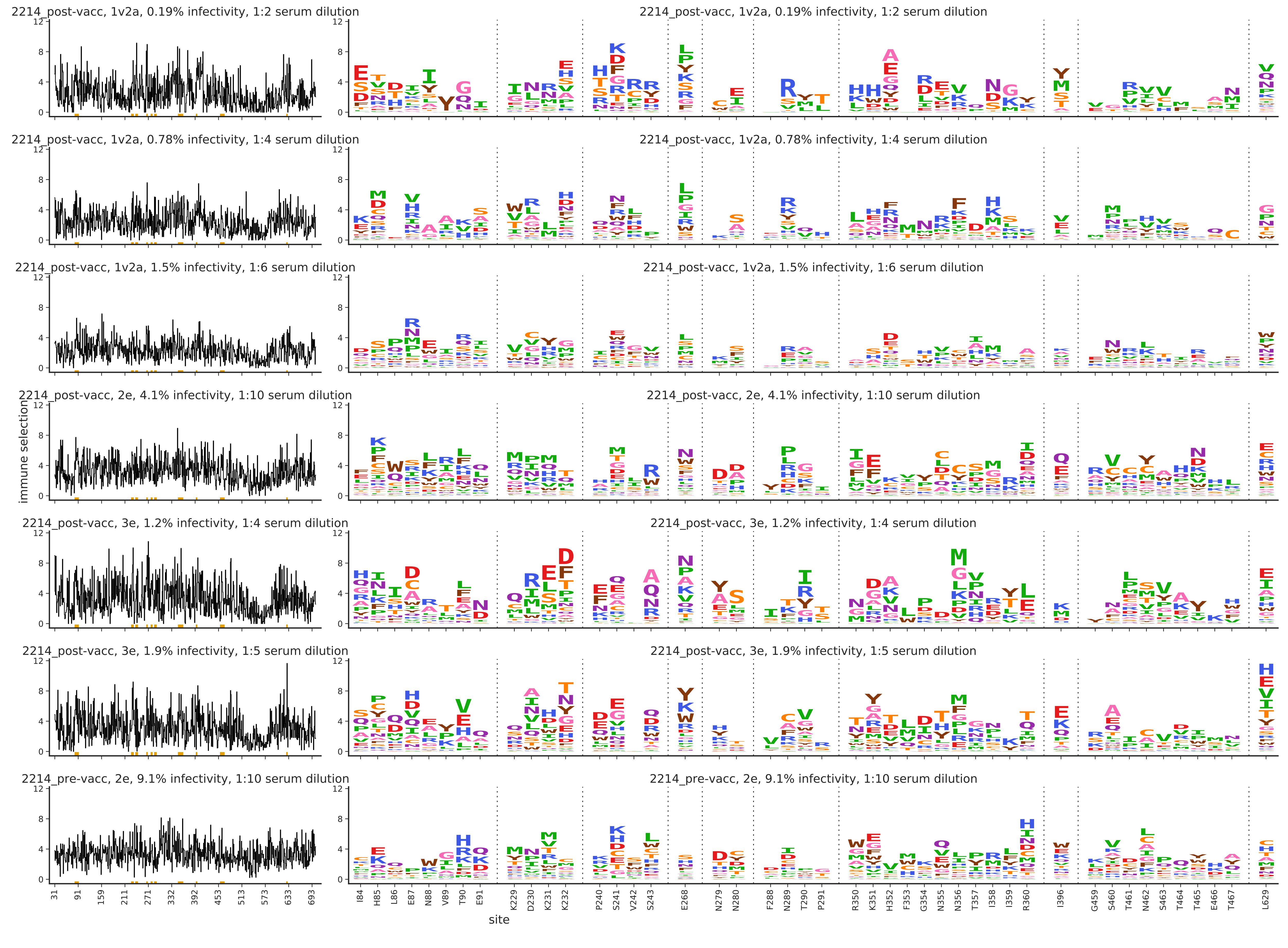

### 2423.pdf

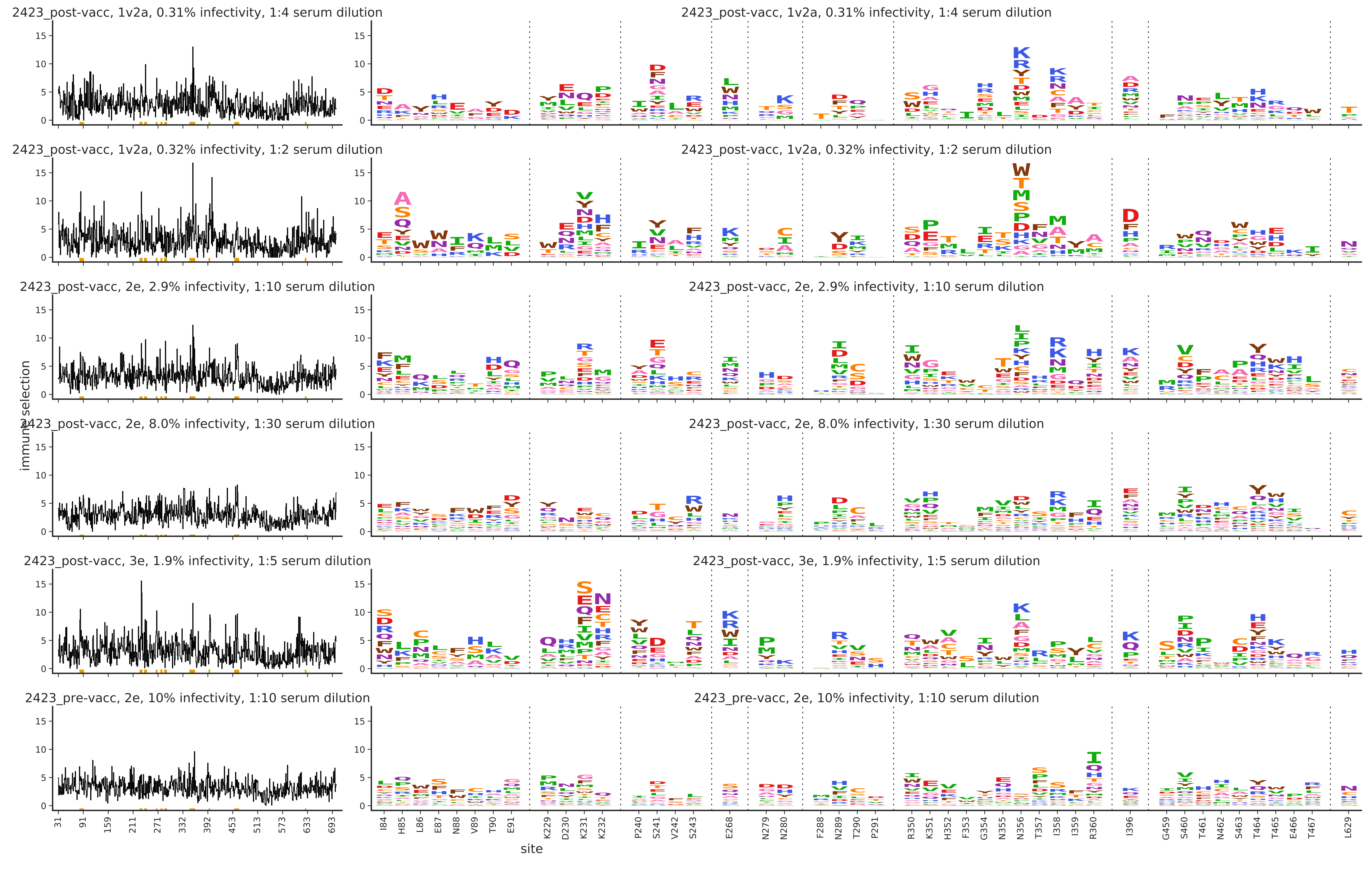

### 2425.pdf

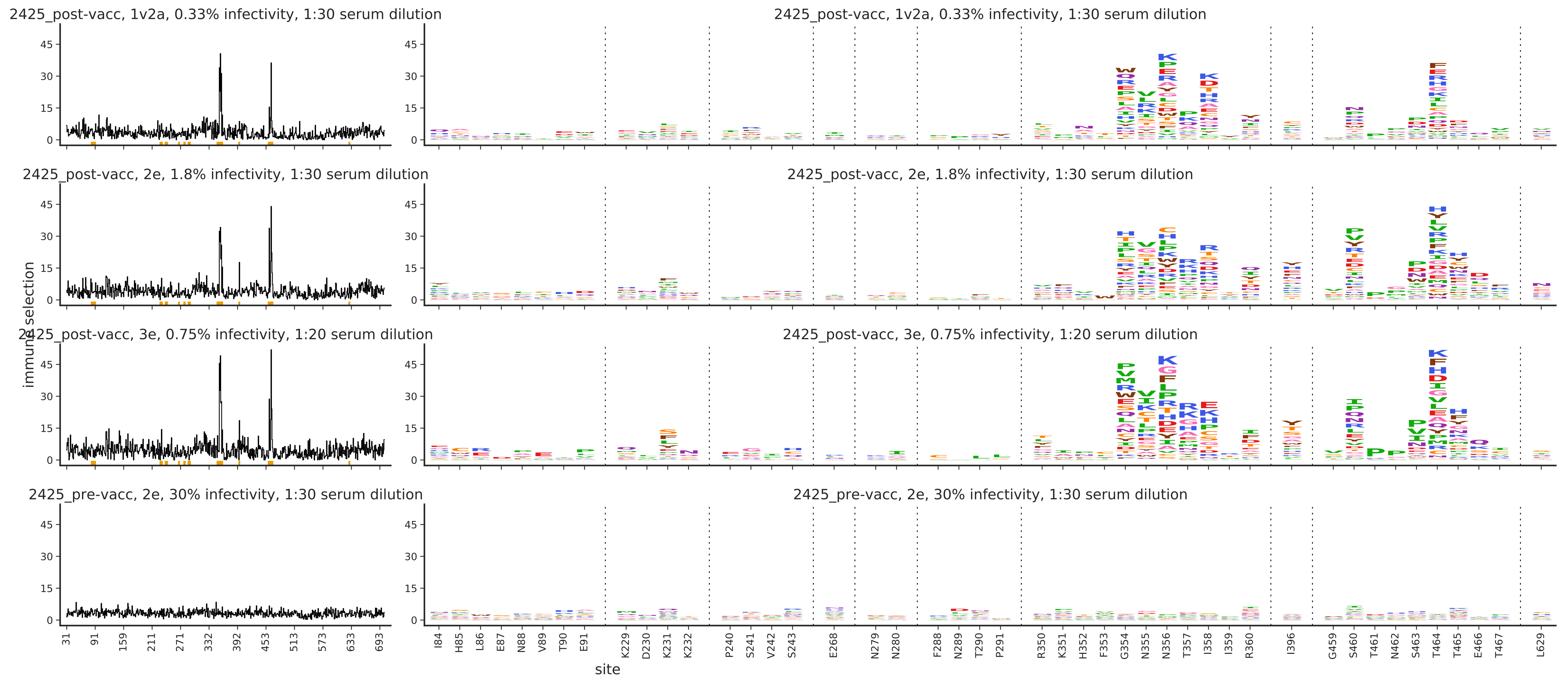

### 5724.pdf

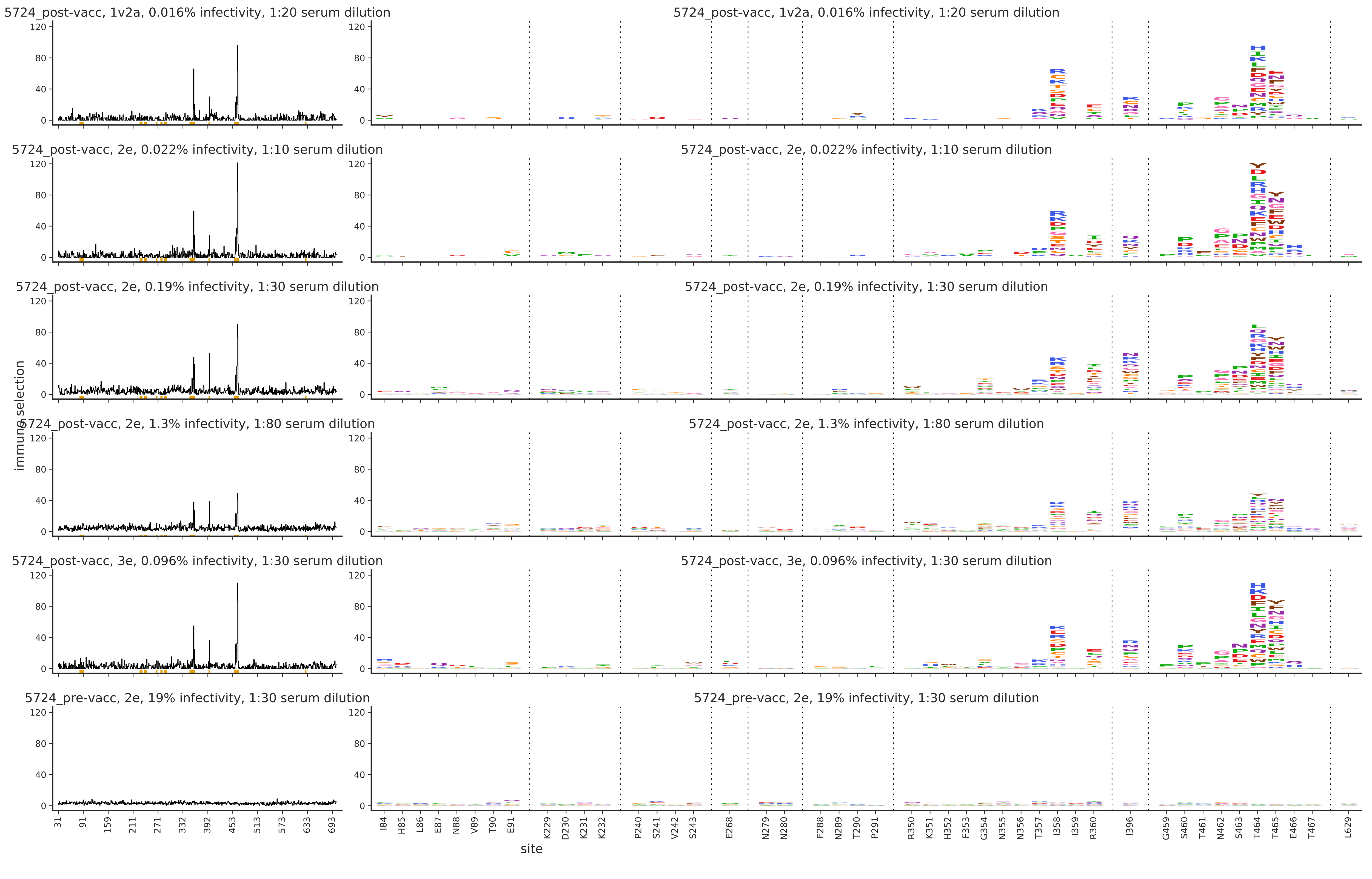

### 5727.pdf

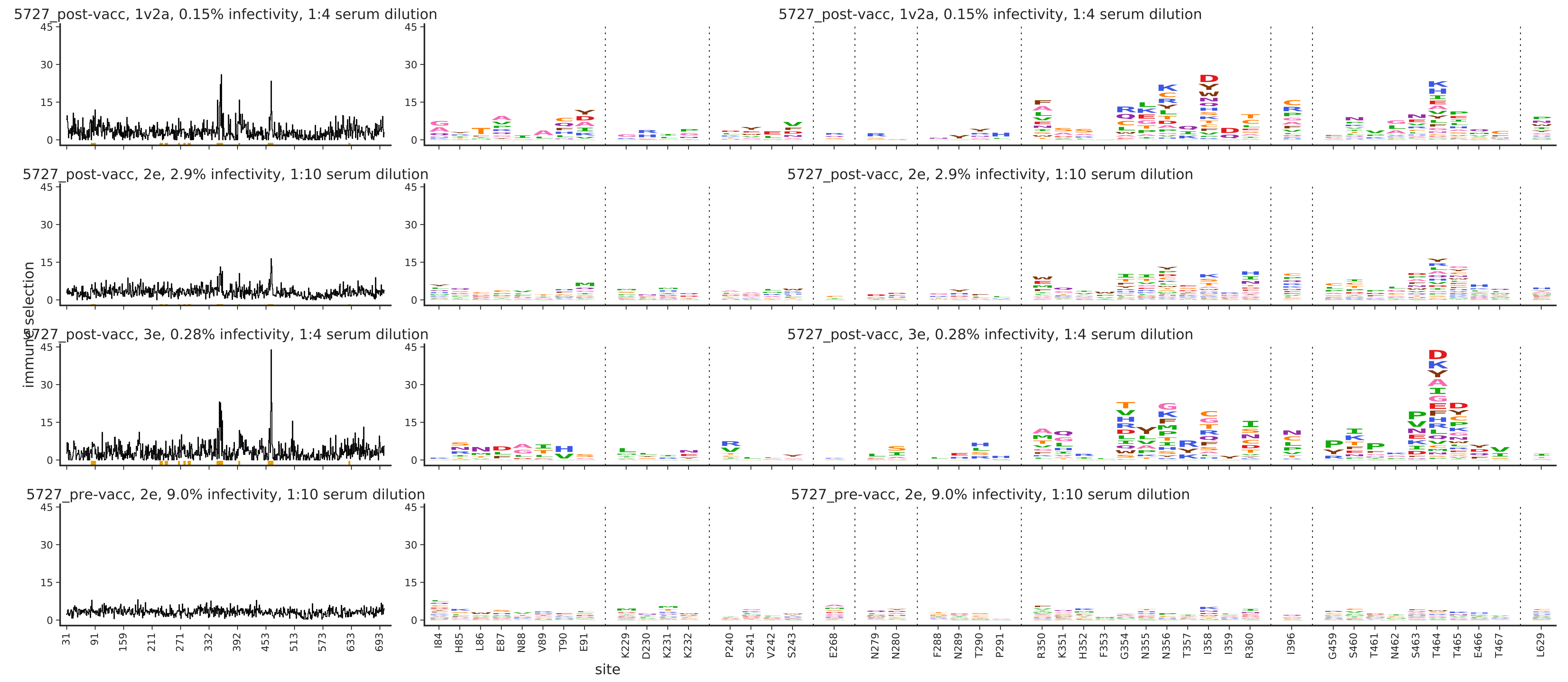

### median-2124-Wk0_diffsel.pdf

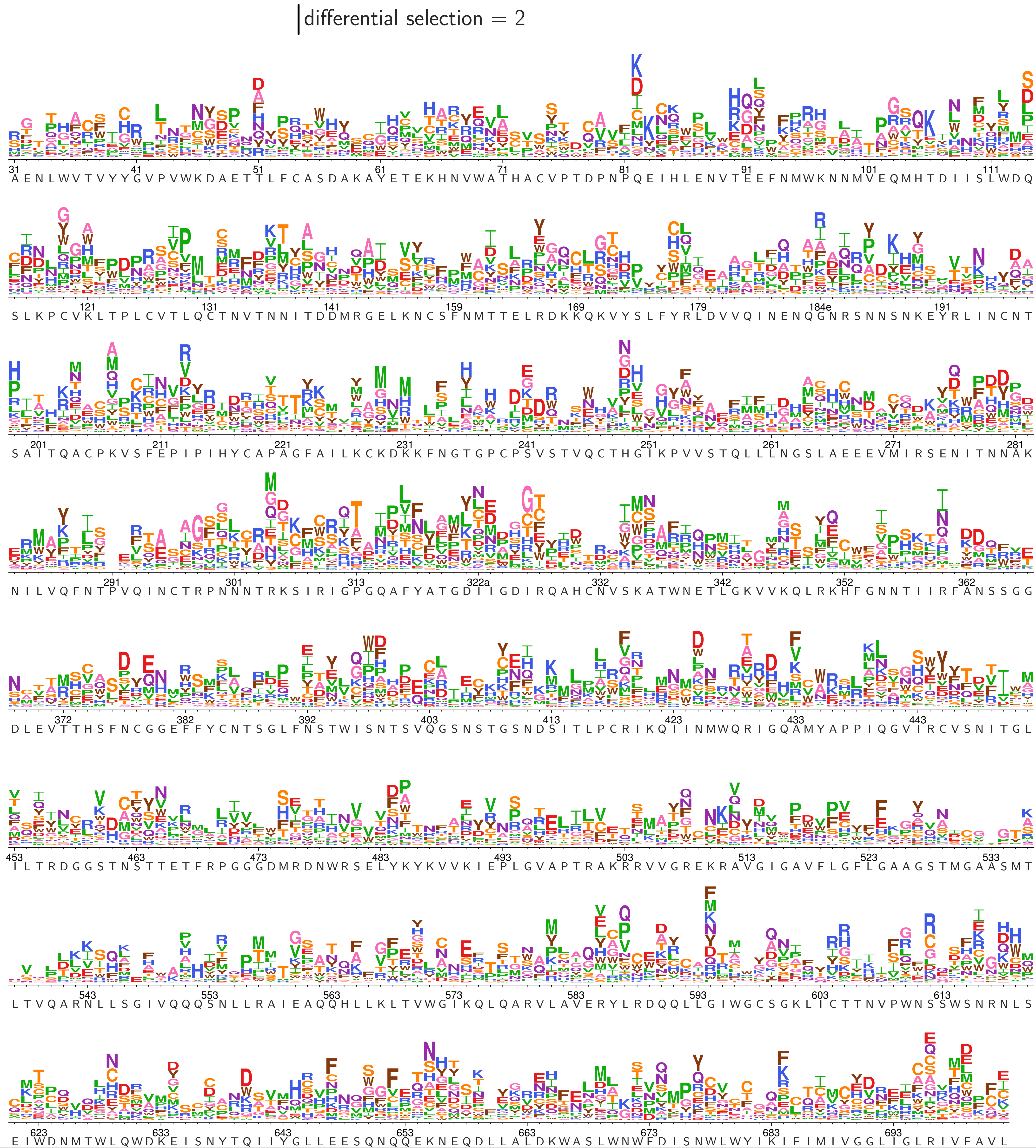

### median-2124-Wk22_diffsel.pdf

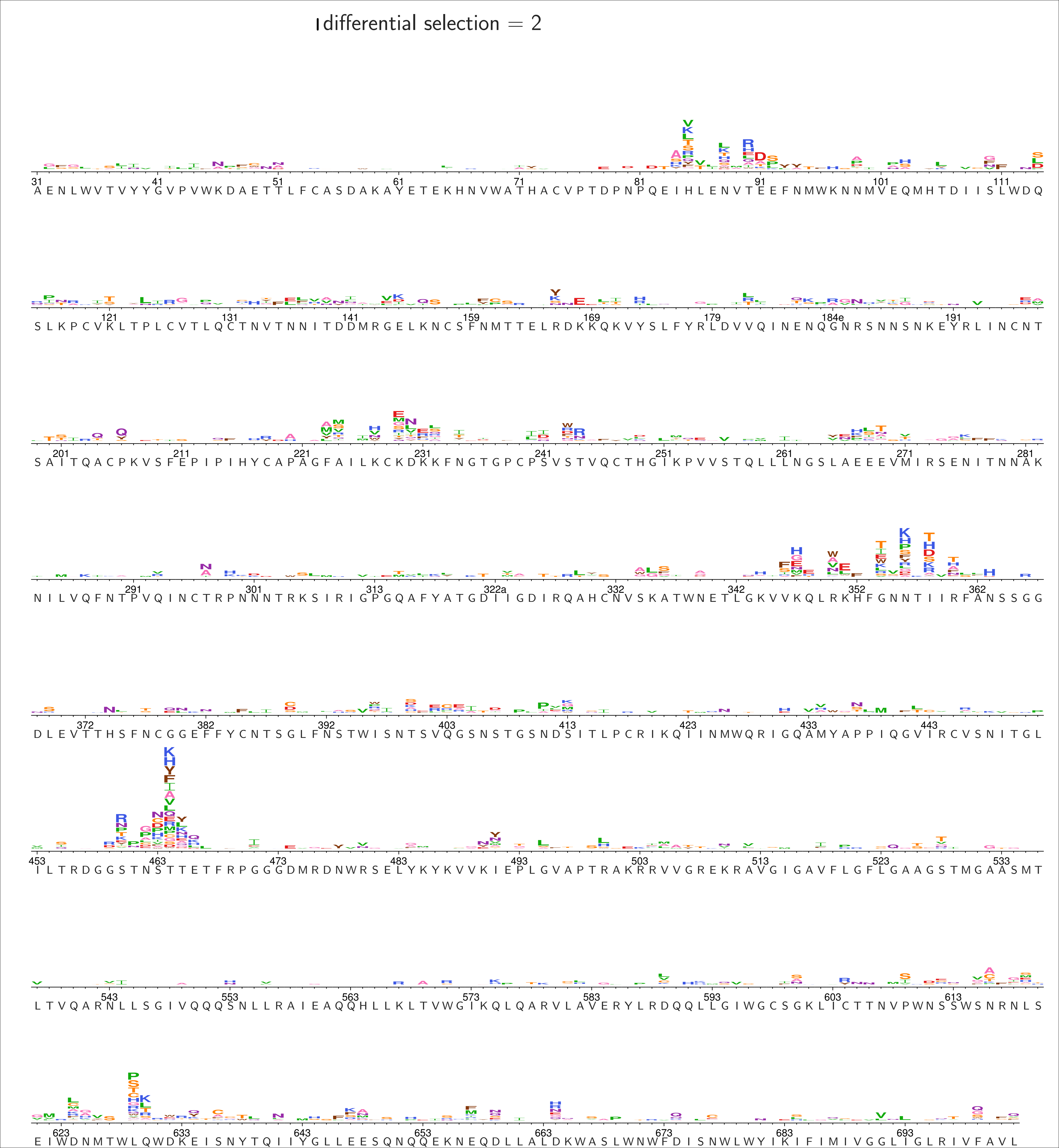

### median-2214-Wk0_diffsel.pdf

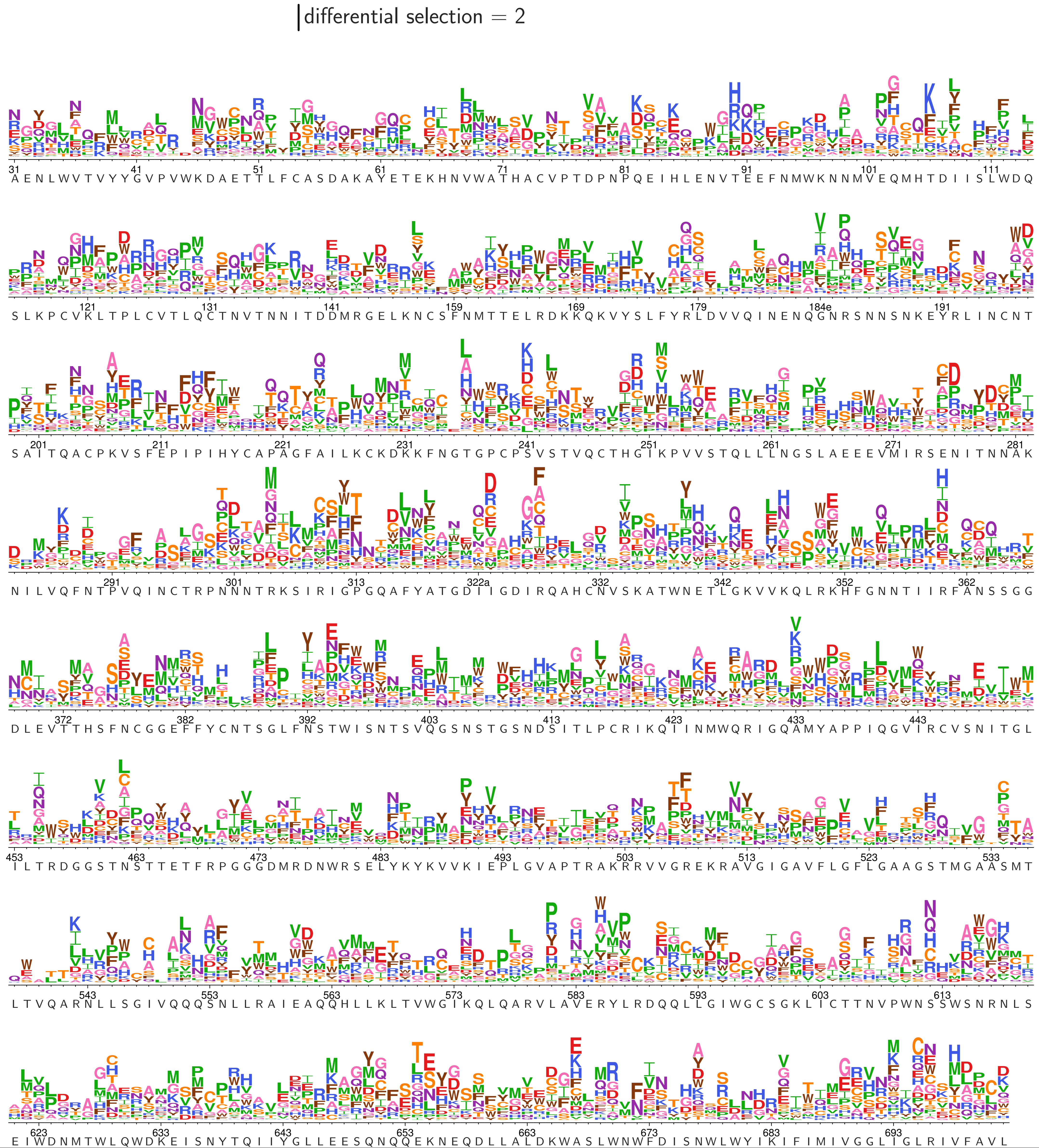

### median-2214-Wk43_diffsel.pdf

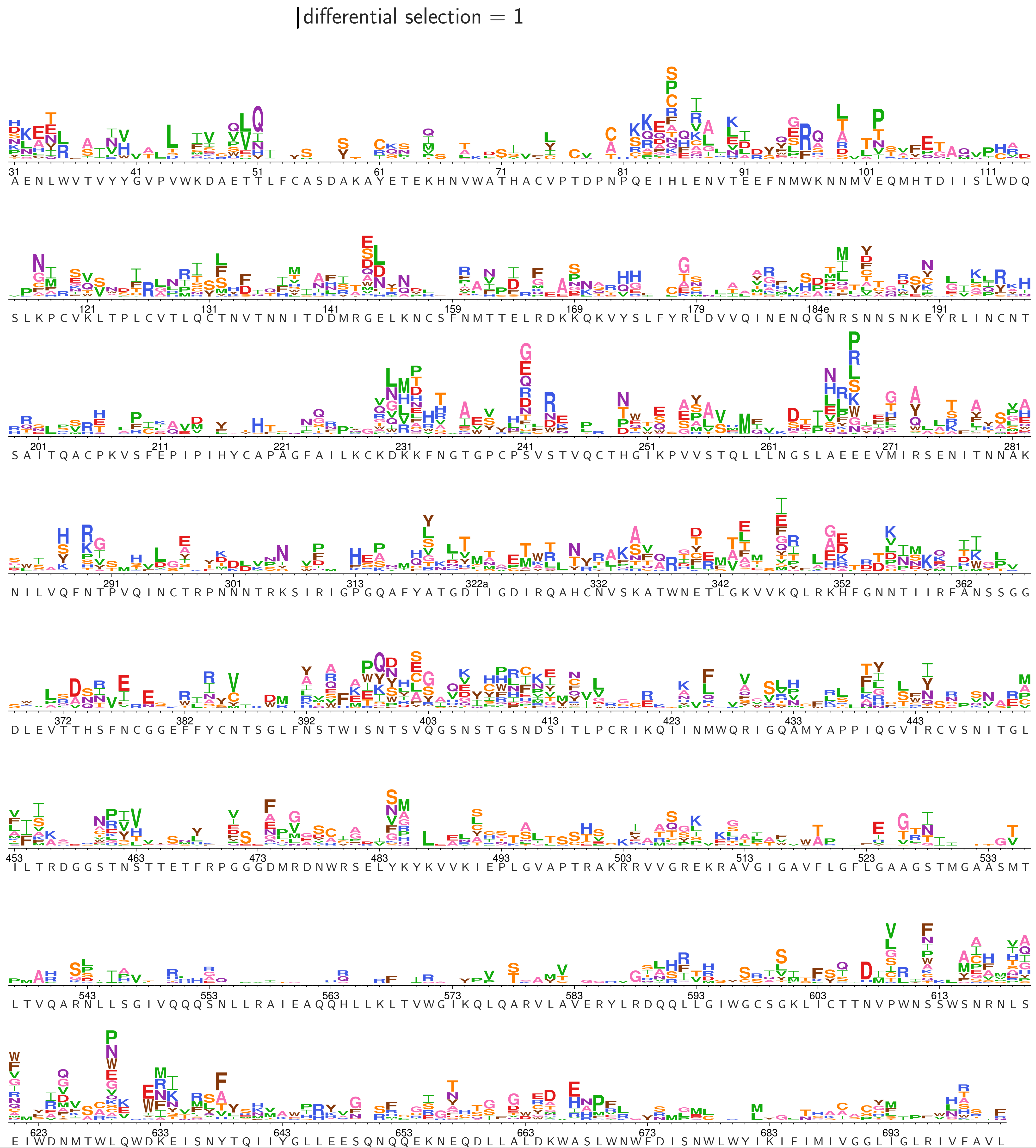

### median-2423-Wk0_diffsel.pdf

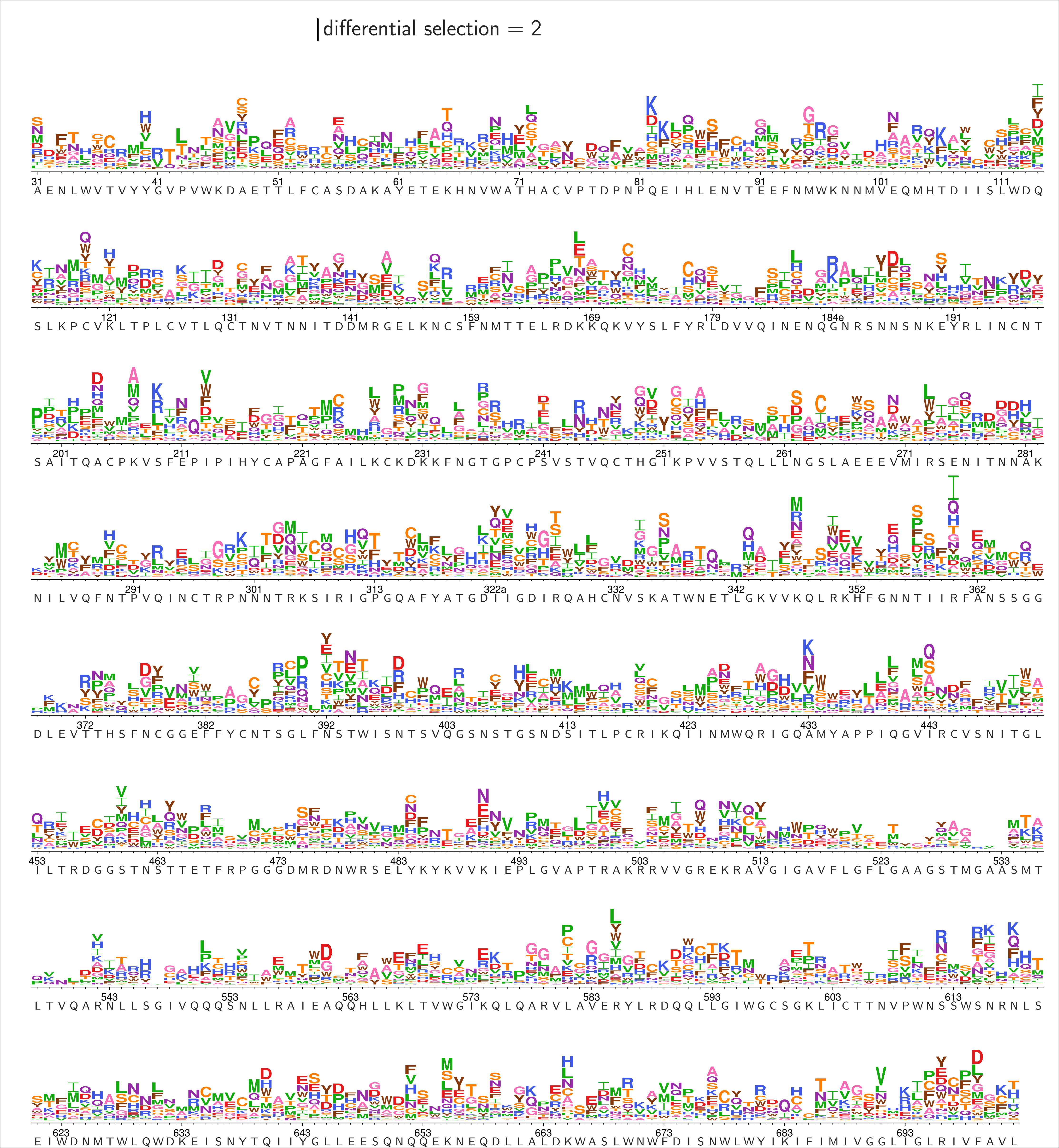

### median-2423-Wk18_diffsel.pdf

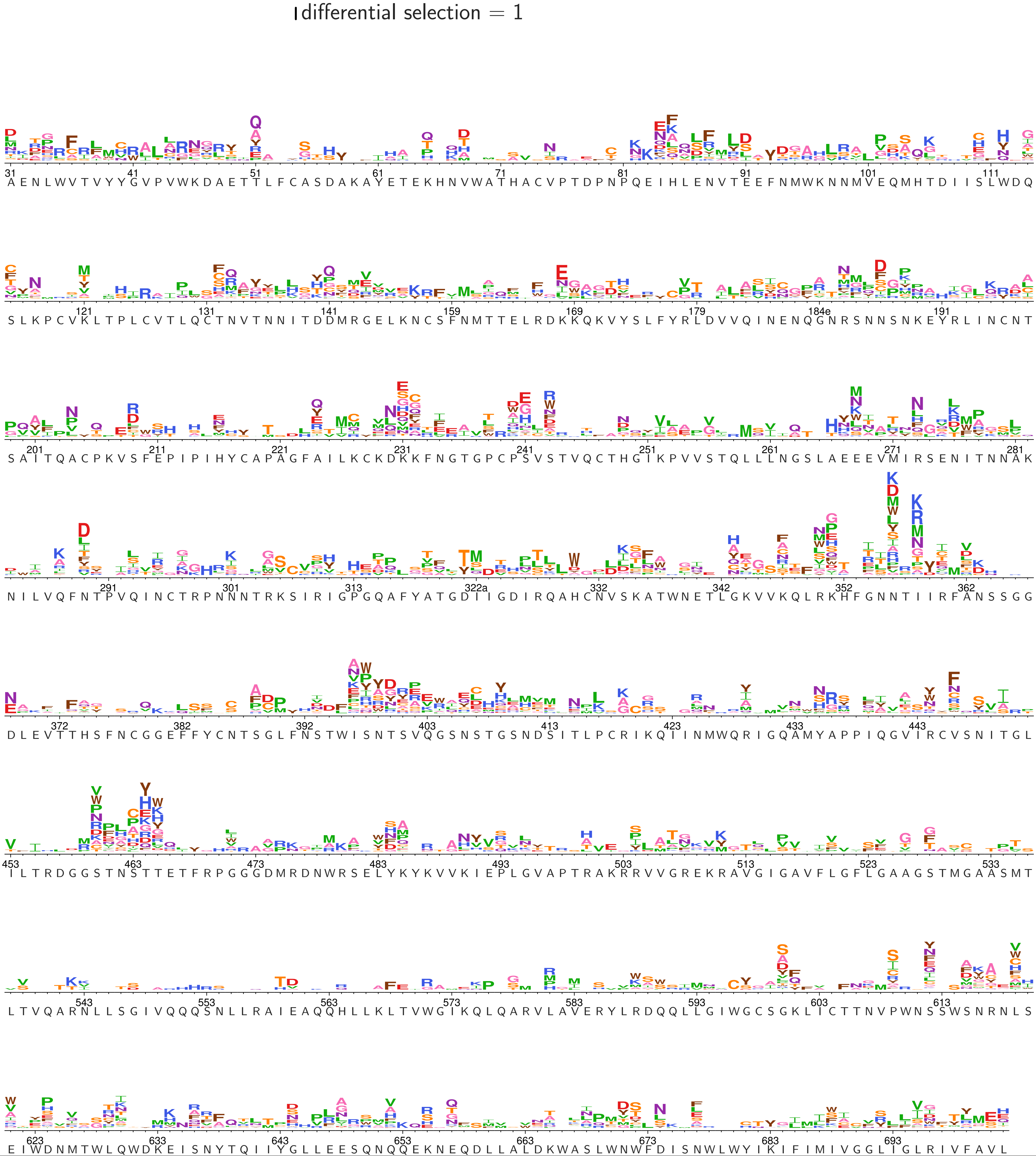

### median-2425-Wk0_diffsel.pdf

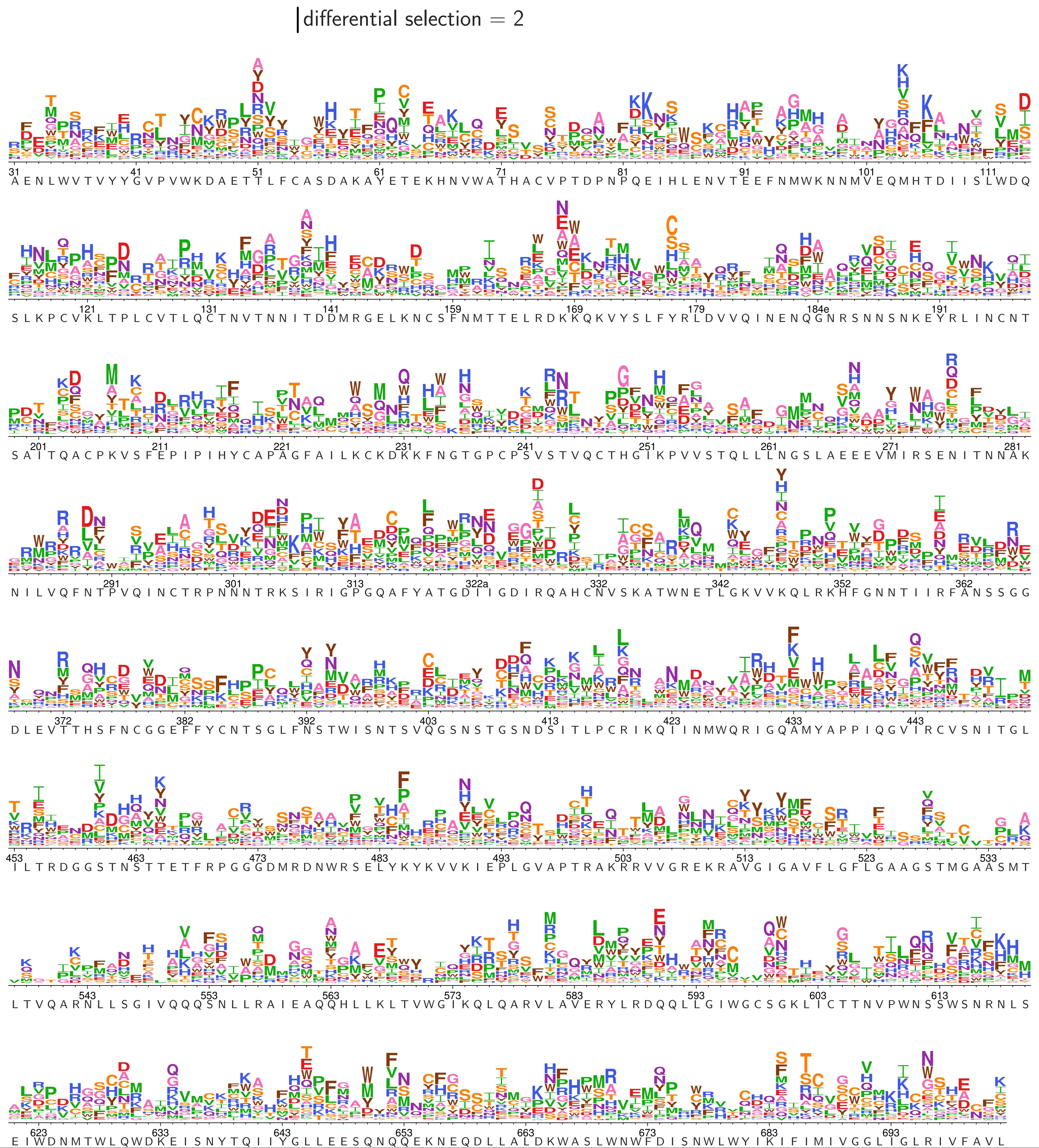

### median-2425-Wk18_diffsel.pdf

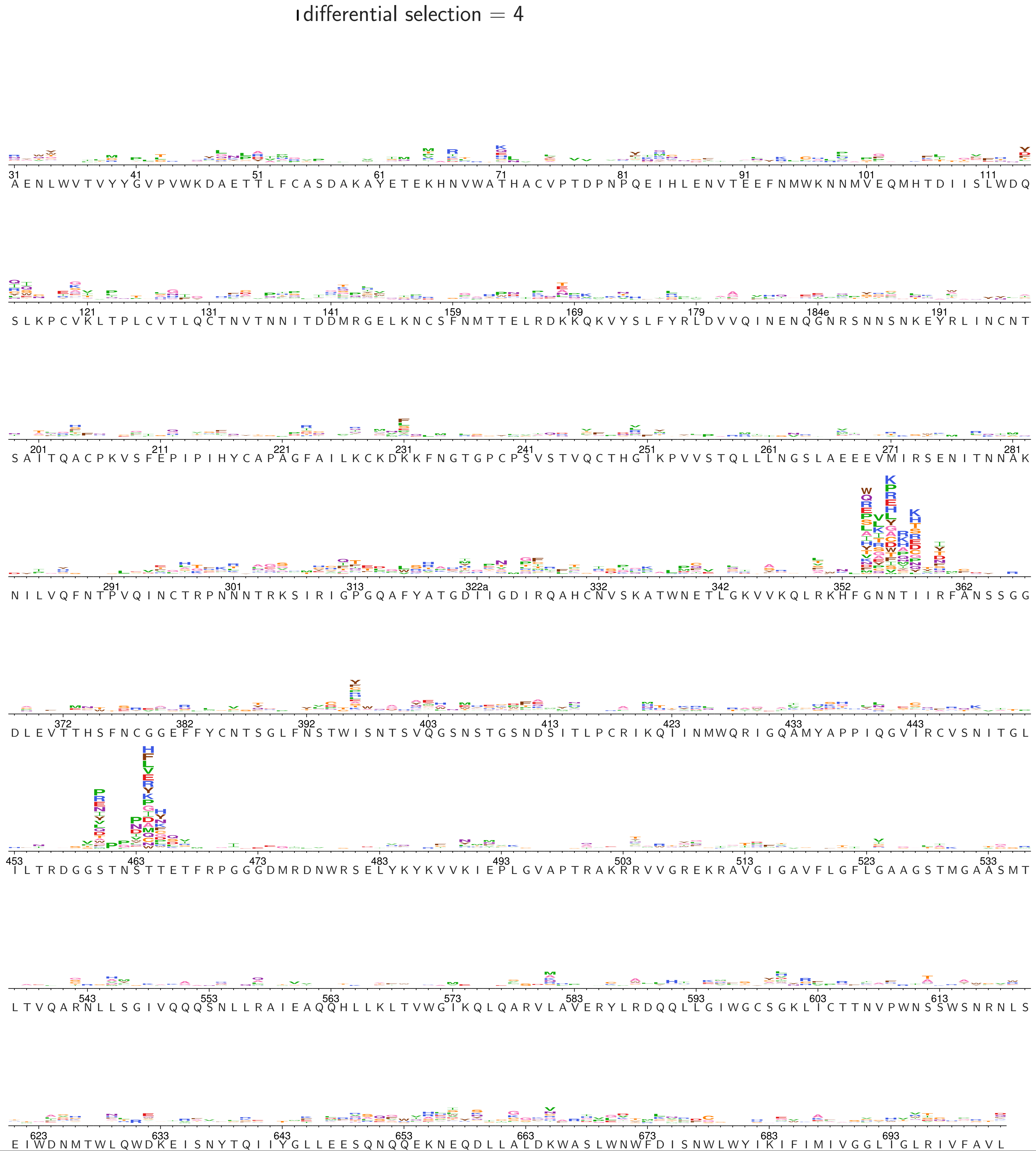

### median-5724-Wk0_diffsel.pdf

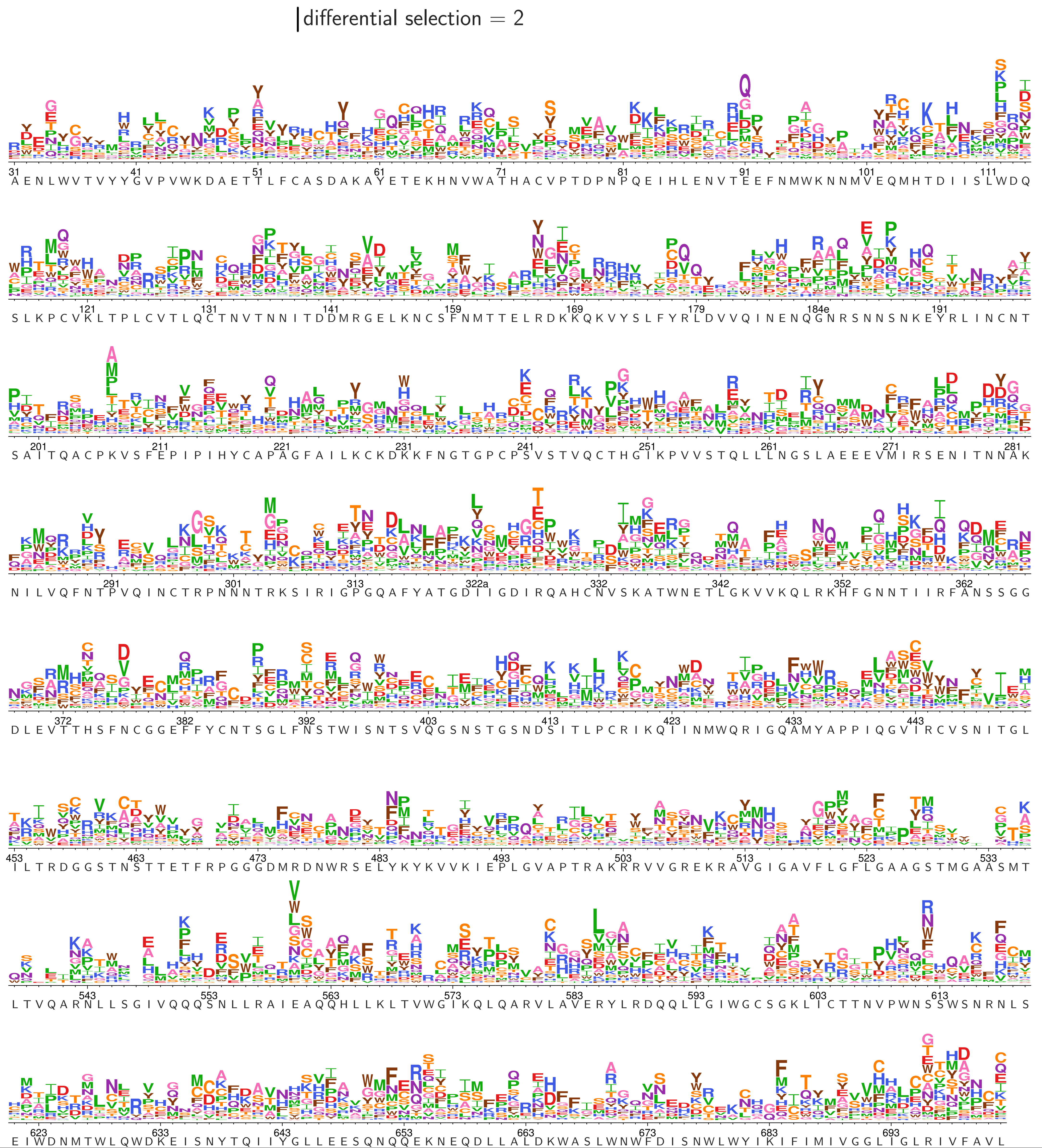

### median-5724-Wk26_diffsel.pdf

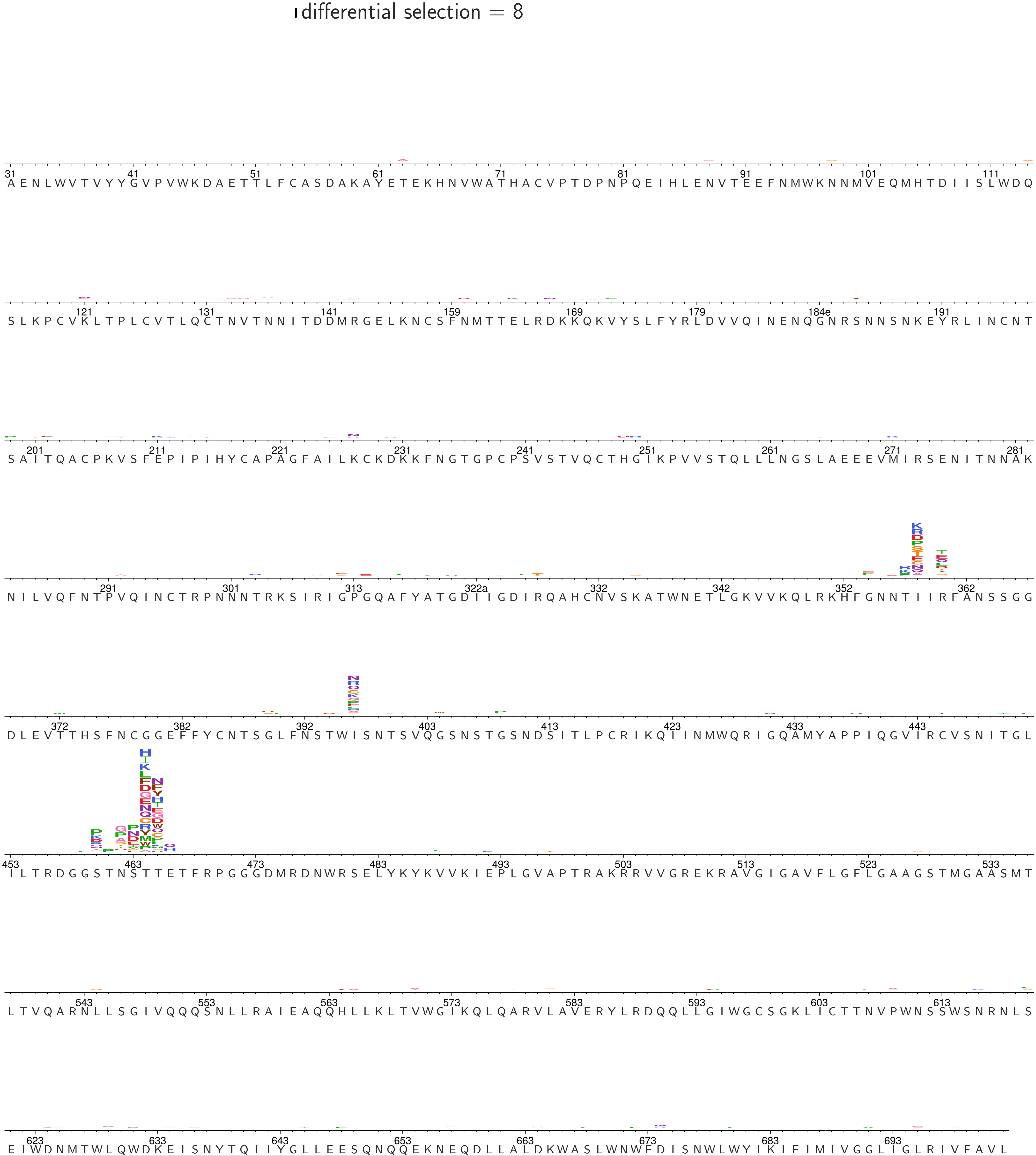

### median-5727-Wk0_diffsel.pdf

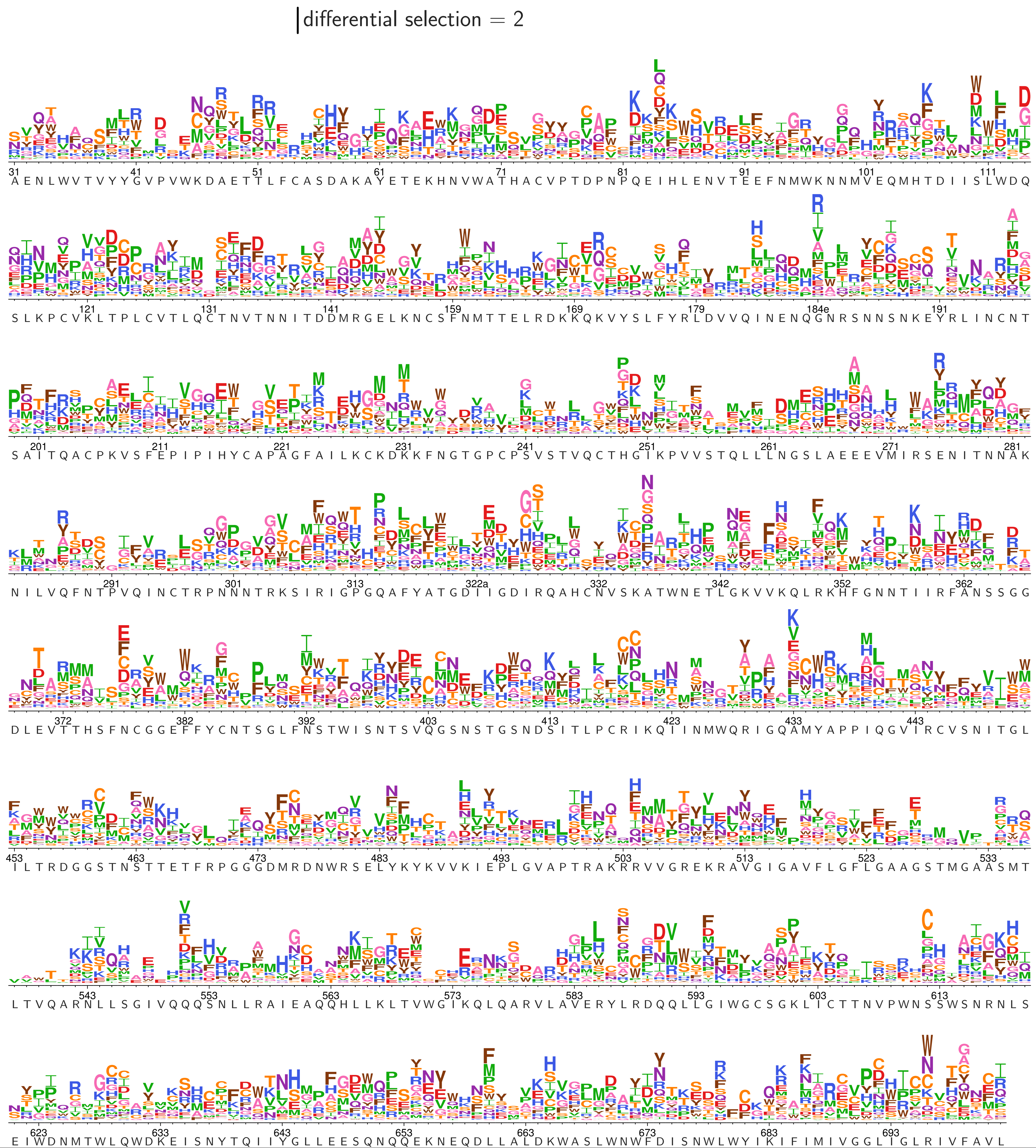

### median-5727-Wk26_diffsel.pdf

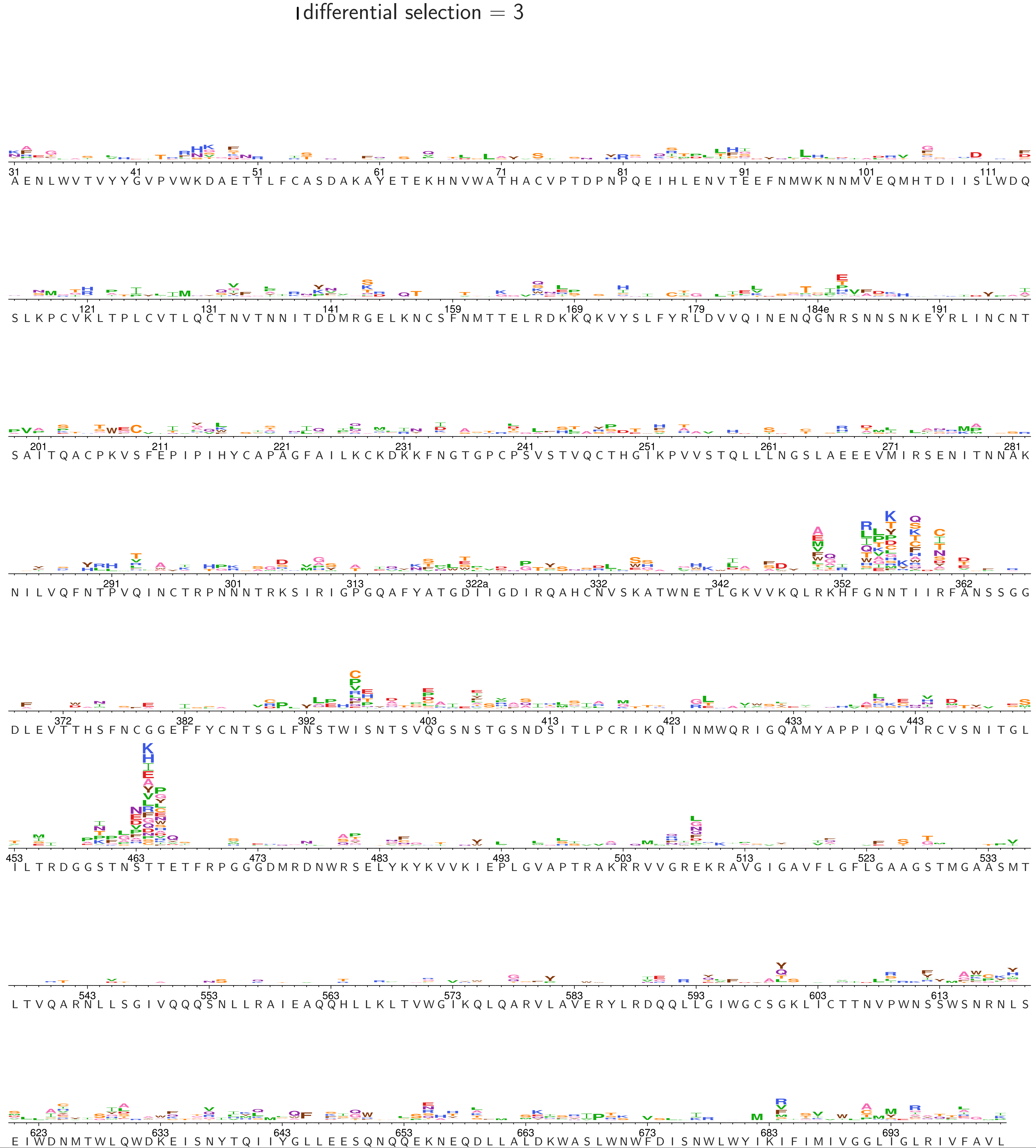
